# Integrated characterization of subsurface media from locations up- and down-gradient of a uranium-contaminated aquifer

**DOI:** 10.1101/712562

**Authors:** Ji-Won Moon, Charles J. Paradis, Dominique C. Joyner, Frederick von Netzer, Erica L. Majumder, Emma R. Dixon, Mircea Podar, Xiaoxuan Ge, Peter J. Walian, Heidi J. Smith, Xiaoqin Wu, Grant M. Zane, Kathleen F. Walker, Michael P. Thorgersen, Farris L. Poole, Lauren M. Lui, Benjamin G. Adams, Kara B. De León, Sheridan S. Brewer, Daniel E. Williams, Kenneth A. Lowe, Miguel Rodriguez, Tonia L. Mehlhorn, Susan M. Pfiffner, Romy Chakraborty, Adam P. Arkin, Judy D. Wall, Matthew W. Fields, Michael W. W. Adams, David A. Stahl, Dwayne A. Elias, Terry C. Hazen

## Abstract

The processing of sediment to accurately characterize the spatially-resolved depth profiles of geophysical and geochemical properties along with signatures of microbial density and activity remains a challenge especially in complex contaminated environments. To provide site assessment for a larger study, we processed cores from two sediment boreholes from background and contaminated core sediments and surrounding groundwater from the ENIGMA Field Research Site at the United States Department of Energy (DOE) Oak Ridge Reservation (ORR). We compared fresh core sediments by depth to capture the changes in sediment structure, sediment minerals, biomass, and pore water geochemistry in terms of major and trace elements including contaminants, cations, anions, and organic acids. Soil porewater samples were matched to groundwater level, flow rate, and preferential flows and compared to homogenized groundwater-only samples from neighboring monitoring wells. This environmental systems approach provided detailed site-specific biogeochemical information from the various properties of subsurface media to reveal the influences of solid, liquid, and gas phases. Groundwater analysis of nearby wells only revealed high sulfate and nitrate concentrations while the same analysis using sediment pore water samples with depth was able to suggest areas high in sulfate- and nitrate- reducing bacteria based on their decreased concentration and production of reduced by-products that could not be seen in the groundwater samples. Positive correlations among porewater content, total organic carbon, trace metals and clay minerals revealed a more complicated relationship among contaminant, sediment texture, groundwater table, and biomass. This suggested that groundwater predominantly flowed through preferential paths with high flux and little mixing with water in the interstices of sediment particles, which could impact microbial activity. The abundant clay minerals with high surface area and high water-holding capacity of micro-pores of the fine clay rich layer suggest suppression of nutrient supply to microbes from the surface. The fluctuating capillary interface had high concentrations of Fe and Mn-oxides combined with trace elements including U, Th, Sr, Ba, Cu, and Co. This suggests the mobility of highly toxic elements, sediment structure, and biogeochemical factors are all linked together to impact microbial communities, emphasizing that solid interfaces play an important role in determining the abundance of bacteria in the sediments.

## 1. INTRODUCTION

As the impact of human processes on land is better appreciated, there is an increasing interest in the biogeochemical processes that drive chemical and physical changes in soil and water and transform and differentially mobilize contaminants. The processes that impact soil and water quality are complex. Abiotic processes of water flow and weathering of rock, chemical transformation of metal and organic contaminants, and interchange of gasses with the atmosphere are mediated by diverse biological processes driven by complex populations of microbes that themselves are affected and dispersed by abiotic factors. Environmental monitoring of these processes has classically proceeded by collection of groundwater in drilled wells and online chemical measurement sites that provide information about bulk chemical concentrations, water flow, temperature and other critical parameters. Interpretation of these data are complicated by the nature of collection since they are naturally an integral of the most mobile elements in the soil and water system that can be collected in homogenous catch-wells. This destroys the spatial information, eludes direct information about the attached and immobilized chemistry and biology. This clearly does not account for the physical changes in the subsurface that mediate key processes but has the advantage of being accessible and temporally continuous (Hazen et al., 1991). In recent years, there is a renewed focus on analyzing the spatially-resolved soil and sediment dynamics along with the differentiated communities of microbes and the processes they harbor.

Here we develop a workflow for characterization of such core-samples that delivers information about the spatially resolved chemistries, physical nature of the sediment matrix, and the bulk biotic properties and show how they differ between cores from two boreholes from very different geochemical positions in a contaminated aquifer and compare these to nearby groundwater wells. We demonstrate how this type of analysis can inform planning for more in-depth investigations of biotic-abiotic interactions in the shallow subsurface and determine the key challenges in the use of this information.

We compared samples from both upgradient and contaminated core sediments, pore waters, and groundwater in nearby monitoring wells freshly obtained from the Oak Ridge Reservation (ORR) and Y-12 Complex field sites in Oak Ridge, TN to determine a full spectrum of site-specific biogeochemical assessments.

Microbes abundant in subsurface environments are more commonly found attached to soil or sediment particle surfaces rather than in suspension in water when nutrient levels are low (Griebler and Lueders, 2009). This particle-attached community composition can be considered a biofilm and is composed of relatively abundant anaerobic bacteria and/or microaerophilic organisms due to low-oxygen zones within the particles (Enzien et al., 1994; LaMontagne and Holden, 2003). These organisms are likely to have an effect on the overall biogeochemical makeup in groundwater systems (Hemme et al., 2010). However, the effect of introduced contaminants on the metabolic potential of groundwater microbes are only vaguely understood (Smith et al., 2018; Smith et al., 2012). We believe the local variation in these parameters strongly affect the distribution and activity of microbes in the soil and thus the overall activity of the site is best predicted by understanding this spatial variation. But accurate measurement of the local physical and chemical parameters that affect biological factors is difficult (Christensen et al., 2018).

For this study we used two freshly drilled boreholes, one in a relatively pristine area up-gradient of a source contaminant and one down-gradient of that contaminated area (S-3 ponds) in the Y-12 National Security Complex, respectively (Smith et al., 2015). These areas were selected to have a similar geology and were considered to be geochemically favorable for microbial nitrate- and sulfate-reduction as previously reported (Smith et al., 2015). We focused on geochemical and bulk biomass (cell counts) differences between background and contaminated subsurface matrix including soil, liquid, and gas phases and the biogeochemical porewater properties along the sediment core depths as compared to homogenized groundwater-only samples. Finally, we determined how these integrated properties of subsurface matrix can be interpreted for future studies. This study reveals an enhanced understanding of how the fundamental geochemical properties of solid, aqueous, and gas matrix, including groundwater flow affect the geochemistry.

## 2. MATERIALS AND METHODS

Here we demonstrate an environmental systems approach for subsurface investigation using the sectioned core sediments and porewater in detail, compared to bulk homogenized results from adjacent monitoring wells.

### 2.1. Study site

The study site is located in the Y-12 National Security Complex in Bear Creek Valley (BCV) in Oak Ridge, Tennessee, USA. The study was conducted within the vicinity of the former S-3 ponds, which contain contaminated waste associated with the refinement of uranium ore (Fig. 1). The geological and hydrological setting within the vicinity of the former S-3 ponds has been previously summarized (Watson et al., 2004). In brief, bedrock depth is 5-10 meters below ground surface (mbgs). Overlying the bedrock are unconsolidated materials comprised of native and non-native porous media including residuum and saprolite while non-native porous media includes alluvium and colluvium which are collectively classified as silty and clayey. The depth to groundwater is ∼2.5-3 mbgs and the water table level is largely controlled by precipitation and infiltration, varying up to 1 meter during major precipitation or drought events. Groundwater flow within the unconsolidated materials is largely controlled by ground surface elevation and is generally south to southwest and towards Bear Creek (Fig. 1).

**Figure 1.**
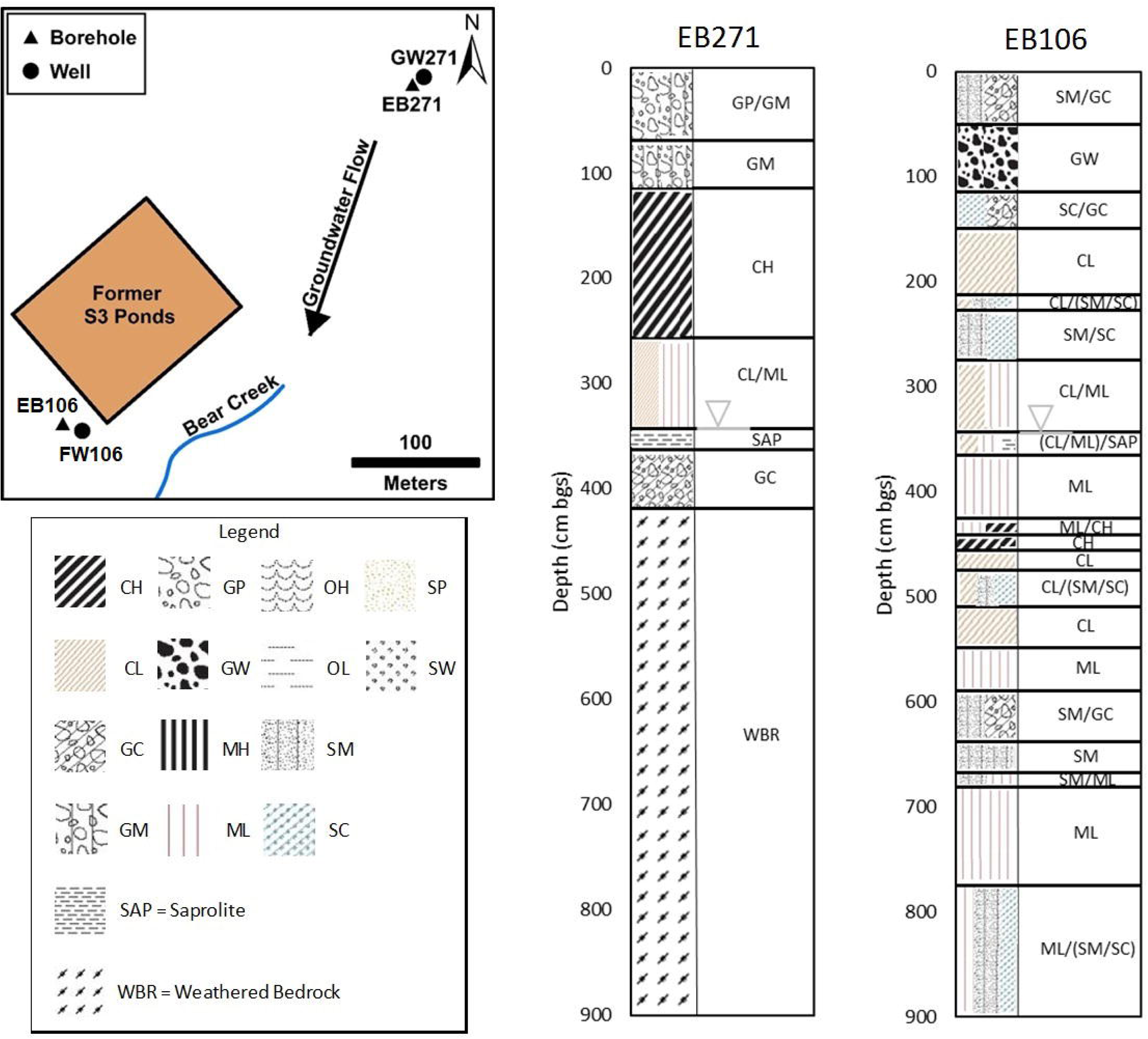
Study site map showing locations of boreholes, wells, former S3 ponds, Bear Creek, and approximate direction of groundwater flow. The unified soil classification system (USCS) of core sediments from borehole EB271 and EB106 based on in-situ visual and manual soil and sediment classification, inverted triangles indicate water table.

We focused on the comparative site-specific core assessment with discrete depth profiles from 22.86 cm core segments and assessed the accompanying subsurface properties using a new borehole in a upgradient area (EB271) and one in a contaminated area (EB106). Both core samples were immediately processed (sampling, handling, and shipping for further simultaneous analyses at various institutions under anaerobic or minimized air-exposed conditions). We performed depth-specific site assessment including sediment minerals and major transition metals, radionuclide concentrations, biomass (total cell counts) of solid media, pH, conductivity, anions, organic acids, and cations. Sediment water characteristics were compared to the same features obtained from groundwater (GW271 and FW106) which likely represents a homogenized mix regardless of the heterogeneity within vadose, capillary, and saturated zones (Supplemental Table 1).

### 2.2. Sediment collection

Core samples of unconsolidated materials were obtained from boreholes EB271 and EB106 (Fig. 1). EB271 is located up-gradient of the former S-3 ponds, whereas EB106 was located down-gradient 21.1 m downstream from the southern corner of the S-3 pond and as such is contaminated with uranium and nitric acid whereas EB271 was not. The boreholes were advanced using a dual tube (DT22) direct-push Geoprobe drill rig resulting in a sealed casing through which undisturbed sediment samples are recovered. Sediment samples were encased in disposable thin-walled polyvinyl chloride (PVC) liners (91.44 cm length x 5.1 cm I.D.) attached to 3.18 cm-outside diameter inner rods (Geoprobe, 2011). Core sediment samples were field-logged for lithological content using the visual-manual Unified Soil Classification System (USCS) to estimate moisture content, density, color, grain size distribution, and other notable field observations. After drilling was completed, the boreholes were filled with a bentonite and cement slurry to approximately 1 mbgs. From 1 mgbs to ground surface, the boreholes were filled with native material. The locations (latitude and longitude) of each borehole were surveyed in the field using GPS (See Supplemental Table 2). Core segments were evaluated for compression or expansion due to deformation during recovery (See Supplemental Tables 3 &4).

Core segments (91.44 cm) were immediately cut to 22.86 cm segments, capped and stored under a nitrogen atmosphere at 4°C after collecting an X-ray fluorescence (XRF) sample by aseptic subsampling of material from the end of each segment. Cores were processed within 24 h of collection with aseptic techniques as follows: Core segments were transferred into an anaerobic chamber (5% H_2_/95% N_2_; Coy Laboratories, Ann Arbor, MI), end caps were removed and intact subcore samples were extracted from each end with an autoclaved syringe barrel with the needle adaptor end removed. Sediment along the core length was exposed by longitudinal cuts through the PVC liner with a liner cutter (DT32/DT325, Geoprobe). Intact subcore samples were taken from various locations along the core length with syringe barrels. The remaining sample was homogenized in a mixing tray with sterile plastic liner and chopped into < 5 mm-sized sediment aggregates with autoclaved stainless steel spatulas.

### 2.3. Sediment analyses

All cores were analyzed for radioactivity and U by Geiger counter and XRF respectively. Sediments were placed into double open-ended XRF sample cups (22.9 mm height × 30.7 mm O.D.) with Mylar film (6 μm thickness) covering one end of the cup (Chemplex Industries, Palm city, FL) and measured by an XRF analyzer (XLp-722, Niton) to measure trace metals including Sr, U, Rb, Th, Zn, and Cu, based on the internal calibration to NIST standard soil sample #2710. The Th reading was normalized to reference material data. Samples were also analyzed by a vacuum XRF (Tracer III-SD, Bruker) to measure light elements and readings were normalized to the previous NIST standard material data.

Mineralogical analyses were performed on samples selected for gas analysis. Only a subset of the samples, regardless of contamination, were classified as “Unrestricted use soil I”, meaning they did not exceed the maximum of 91 pCi/g, and were allowed to be analyzed. For this subset, the remaining homogenized sediments were centrifuged for pore water (as below) and were then dried in a fume hood at room temperature, ground and loaded into sample holders. The tubes were covered with Kapton film and adhesive as part of the radiological safety protocols, but the film itself interfered with the low angle range for clay mineral identification. The mineral assemblage was confirmed with an X-ray diffractometer (XRD, X’pert PRO, PANalytical, Natick, MA) (Moon et al., 2014) equipped with Mo-Kα radiation at 55 kV/40 mA between 3-25° 2θ with 0.55° 2θ/min. A semi-quantification of mineral constituents was performed by interlocked ratios (Moon et al., 2000) by both the reference intensity ratio (Chung, 1974) and mineral intensity factors (Ahn, 1992).

Inductively coupled plasma mass spectrometry (ICP-MS) samples for trace elemental analysis were collected and analyzed as previously described (Ge et al., 2019). In short, three approximately one gram samples of homogenized sediment were sampled from each segment. The moisture content and dry mass of samples were determined. Then the samples were microwave digested in 10 mL of concentrated nitric acid as described in EPA Method 3051A (EPA, 2007). After microwave digestion, samples were diluted for ICP-MS analysis, resulting a final dilution of 1:5 or 1:50 in 2% (vol/vol) nitric acid. Fifty-three elements were analyzed by ICP-MS (Supp Data #1). Where indicated, the three environmental replicates where averaged and reported with standard deviations (SD).

### 2.4. Porewater and groundwater collection

The sediment pore water was collected by a drainage centrifuge method (De Goffau et al., 2012) from 20 g of homogenized sediment in each core segment in an anaerobic glove bag. A filter device (Amicon Ultra 100K MWCO, Merck Millipore, Berlington, MA) was used to centrifuge wet core sediments (3,000 x g, 4°C, 1 h). When processing a large number of samples, this less laborious and cheaper method is preferred for extracting pore water and to determine its solute concentration (De Goffau et al., 2012). Since the extracted sediment pore water volumes were relatively small, samples were transferred into GC glass vials with screw caps (PTFE/red silicon septa) in an anaerobic glove bag. Sulfide analysis was immediately initiated to measure dissolved total sulfides.

Groundwater samples were collected from neighboring monitoring wells, GW271 for EB271 and FW106 for EB106, respectively (Fig. 1) by low-flow purge and sampling (Smith et al., 2015). Groundwater was purged with a peristaltic pump connected to dedicated down-well tubing installed at the mid-screen of the wells. A water quality indicator probe (Troll 9500, In-Situ Inc., Fort Collins, CO) was connected in-line to a flow-through cell to monitor pH, conductivity, and oxidation-reduction values during purging. Once these parameters stabilized, samples were collected in an appropriate aseptic/autoclaved container, preserved, and transported for laboratory analysis.

### 2.5. Porewater and groundwater analysis

Dissolved total sulfide was determined using a HACH spectrophotometer (DR2800, HACH Method 8131) and an UV-spectrometer (HP8453, Hewlett Packard, Palo Alto, CA). Samples were analyzed against prepared standards (5-800 μg/L) in 1.5 ml semi-micro cuvettes (Brand, Germany). To determine anions and organic acids, filtered (0.22 μm pore-sized filters) pore waters were diluted into 1.5 ml vials. Concentrations of anions (fluoride, chloride, nitrite, bromide, nitrate, sulfate, and phosphate; working range of 0.1-200 mg/l) and organic acids (lactate, acetate, propionate, formate, pyruvate, butyrate, succinate, oxalate, fumarate, and citric acid; working range of 5-200 μM) were determined on a Dionex™ ICS 5000^+^ series with Dual Pump, Dual Column system (ThermoFisher Scientific, Waltham, MA) equipped with an AS11HC column at 35°C with a KOH effluent gradient of 0-60 mM at 1.3 ml/min.

The cations (lithium, sodium, ammonium, potassium, magnesium, and calcium; working range of 5 μg/l-5 00 mg/l) were determined with a Dionex™ ICS 5000^+^ series with a CS12-A column at 35°C with an isocratic 20 mM methanesulfonic acid effluent at 1 ml/min. Samples were acidified of 10 vol.% 1M HCl solution before analysis.

Sediment pH was measured with a combination of a pH electrode (8103 Ross probe, ThermoOrion) and an Orion EA-920 Expandable ion analyzer (ThermoOrion, Beverly, MA). Dried core sediments in the XRF sample holder were rehydrated with 10 mM CaCl_2_ solution (sediment:solution ratio of 1:2 kg/l) equilibrated for 60 min and centrifuged (3,000 x g, 5 min, 4°C). Conductivity was also measured on the same samples with a combination of a conductivity cell (Orion 011010) and a conductivity/TDS/salinity meter (Thermo Orion 115A^+^). The blank 10 mM CaCl_2_ solution exhibited 4.46 mS/c.

### 2.6. Sediment gas analysis

Homogenized core sediment (2 g) from selected segments of the unsaturated (vadose), capillary fringe, and saturated zones was placed in a 10 ml serum bottle in the anaerobic glove bag and sealed with a blue butyl rubber stopper and aluminum crimp and then stored in the dark for 1 week. Unfiltered groundwater (2 ml) from the closest neighboring monitoring well (GW271 for EB271 and FW106 for EB106) was injected into the serum bottle by syringe and needle immediately after groundwater collection. Those serum bottles including sediment and groundwater were mixed by shaking in the end-to-end shaker (3 h) and aluminum foil-covered samples were stored in the glove bag for 4 weeks. Headspace gases from the incubations (2.5 mL) were transferred to a 12 ml exetainer (Labco, Lampeter, UK) and CH_4_, CO_2_, and N_2_O were measured by gas chromatography (Model 8610, SRI Instrument, Torrance, CA) with nitrogen as carrier, a 182.9 cm HayeSep D column (SRI Instrument) and TCD, ECD and FID detectors at University of Washington.

### 2.7. Biomass analysis

Direct cell counting was performed by a flocculation technique (Sinclair and Ghiorse, 1989). Bacterial cells were released from sediment particles in a solution of CaCl_2_, Tween and sodium pyrophosphate with sonication and vortexing. Sediments were centrifuged (3000 x g, 3 min, 4°C) and the supernatant was filtered (0.2 μm pore-sized black polycarbonate membrane) to concentrate the cells. The cells were stained for 2 min in the dark with 0.25 μM acridine orange and unbound stain rinsed from cells with phosphate buffered saline. Stained cells were viewed with epifluorescence microscope (Zeiss Axioskop, Germany) with a fluorescein isothiocyanate filter. Microbial biomass C in core sediment sample was measured using chloroform fumigation extraction method. Each sediment sample was divided into two subgroups. For fumigation group, 5 g fresh sediment was measured into a 50-ml centrifuge tube. A cotton ball was put 4–5 cm above the sediment sample. Then 2.5 ml of CHCl_3_ was added to the cotton ball. The tube was stored in dark at room temperature for 7 days. At day 3, another 2.5 ml of CHCl_3_ was added to the cotton ball. After 7 days, chloroform was evacuated in a fume hood and sediment sample was extracted with 20 ml of K_2_SO_4_ (0.5 M) on a shaker for 1 hr. The extracts were filtered through a qualitative filter into a plastic scintillation bottle, then stored at 4°C for TOC analysis. For control group, another 5 g fresh sediment was measured into a 50-ml centrifuge tube and then extracted immediately using the same method as fumigation group. The TOC content of extract was measured by TOC-5050A Total Organic Carbon Analyzer (Shimadzu, Japan). Biomass C was calculated based on the TOC difference between fumigation group and control group.

### 2.8. Organic C, Total C, and Total Nitrogen

The sediment samples were oven-dried at 70°C and ground to a fine powder. A representative sample (∼0.5 g) was loaded into a ceramic sample boat and combusted in an oxygen atmosphere in an Elementar Vario Max Analyzer (Elementar Americas Inc, NY). Elemental C and N were finally converted into CO_2_ and N_2_. These gases were then passed through the infrared cells to determine CO_2_ and a thermal conductivity cell to determine N_2_. For organic C measurement, acid-treatment was applied to remove inorganic C prior to combustion.

### 2.9. Micro computerized tomography (Micro-CT) sample preparation and image data collection

Non-homogenized sediment samples were excised from the field cores by inserting 3 cc syringes (in which the front sections with needle adapter were removed to expose the bore of the main cylinder) into the sediment to isolate ∼8 mm diameter plugs. These plugs were transferred into 2 ml Eppendorf microcentrifuge tubes and stored at 4°C. To prepare specimens for micro-CT imaging, Kapton tubes (1.4 mm ID, 0.051 mm wall thickness, Cole-Parmer, Vernon Hills, IL) cut into 4 cm lengths were pressed into the core plugs to take samples of sediment. Tubes loaded in this manner contained an intact plug of sediment 1.4 mm in diameter and ∼ 0.5 cm in length. To preserve moisture, the open ends of the tubes were sealed. Samples were stored at 4°C until analysis.

For analysis, sample tubes were placed vertically into the specimen mount of the micro-CT beamline (Advanced Light Source, beamline 8.3.2), positioned between the source X-ray beam and the detector. In this experimental configuration, X-rays passing through the sample strike a scintillator which in turn is imaged onto a CCD camera (PCO Edge, 2560×2160) with a resulting pixel size of ∼0.7 μm. The tunable X-ray source of the beamline was adjusted for an energy setting of 22 keV. Under these conditions a cylindrical volume of sediment about 1.5 mm in diameter and 1.5 mm in length could be imaged in about 20 minutes. Raw image data were processed and reconstructed into image volumes with the Xi-CAM analysis platform using default parameters. Each resulting dataset is approximately 50 GB in size. Reconstructed image volumes were analyzed, and sectional images prepared with the ImageJ and Avizo software packages.

## 3. RESULTS

### 3.1. Sediment characterization

Boreholes EB271 and EB106 were advanced to ∼5 mbgs and ∼8 mbgs respectively due to a lack of drill advancement. It is likely that consolidated or semi-consolidated material, e.g., weathered bedrock (Fig. 1), was responsible. It should be noted that weathered bedrock was observed in EB271, but not in EB106 (Fig. 1). The first meter of sediments in both boreholes were predominantly gravels and sands; whereas, sediments below this were predominantly silts and clays (Fig. 1). The first meter of sediment was not analyzed due to fire-ant restrictions for shipping. The depth to groundwater in both boreholes was approximately 3.5 mbgs. The depth to weathered bedrock (5 to 8 mbgs), the predominance of silty and clayey sediments, and the depth to groundwater (∼3.5 mbgs) were expected based on previous studies at the site (Watson et al., 2004).

The moisture, relative density, and predominant mean grain size in both boreholes were notably variable with depth (Fig. 2). The moisture of sediments from EB271 showed a sharp increase near the water table followed by a sharp decrease (Fig. 2). This indicated the amount of free water within sediments below the water table varied substantially. EB106 was similarly variable but showed no spike (Fig. 2).

**Figure 2.**
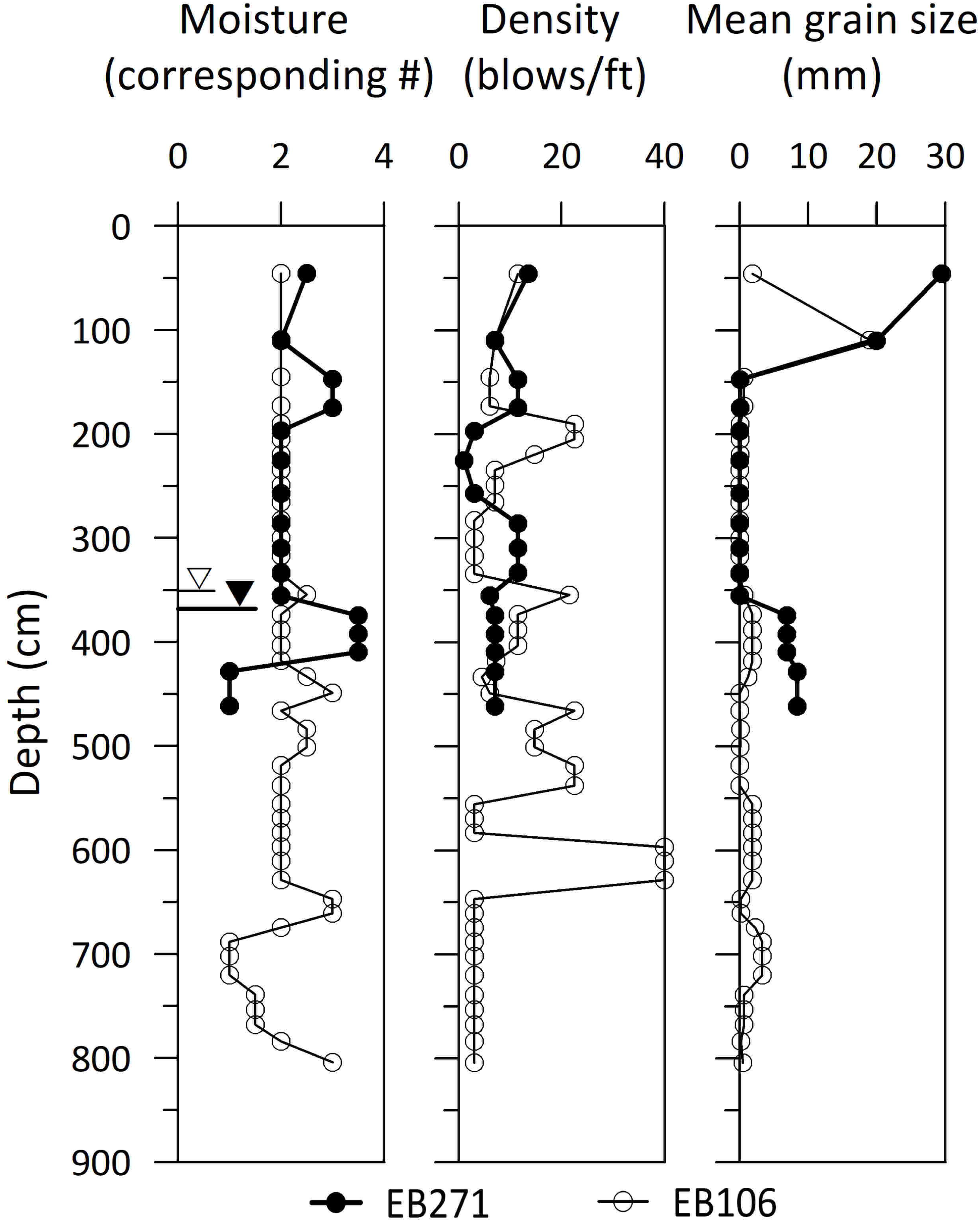
Physical properties (moisture content, density, and mean grain size) of core sediments from background borehole EB271 (closed circles and triangles) and contaminated EB106 (open circles and triangles) based on *in situ* visual and manual soil and sediment classification (USCS). Moisture: 1=dry, 2=moist, 3=wet, 4=saturated. Open and closed triangles show the water table depth.

With respect to trace elements, the prominent difference between the two boreholes was the radioactive material (U and Th) with depth as expected (Fig. 3). Upgradient EB271 sediment contained up to 17.2 mg/kg U and 2.7 mg/kg Th while contaminated EB106 sediment had 249 mg/kg U and 97.5 mg/kg Th, as determined by XRF analysis. The EB271 sediment maintained a low steady concentration of U or Th along the depth to the bed rock layer where coring terminated. In contrast, EB106 sediment exhibited simultaneous U had sharp peaks at 492 cm and 775 cm as well as a broad peak at 183-212 cm bgs in the vadose zone. Th had sharp peaks at 492 cm and 709 cm bgs. U and Th were higher at depths containing clumps of saprolite even though they were mainly composed of clayey silt at the capillary fringe or just below the water table with the lower peak occurring within the less permeable bedrock layer.

**Figure 3.**
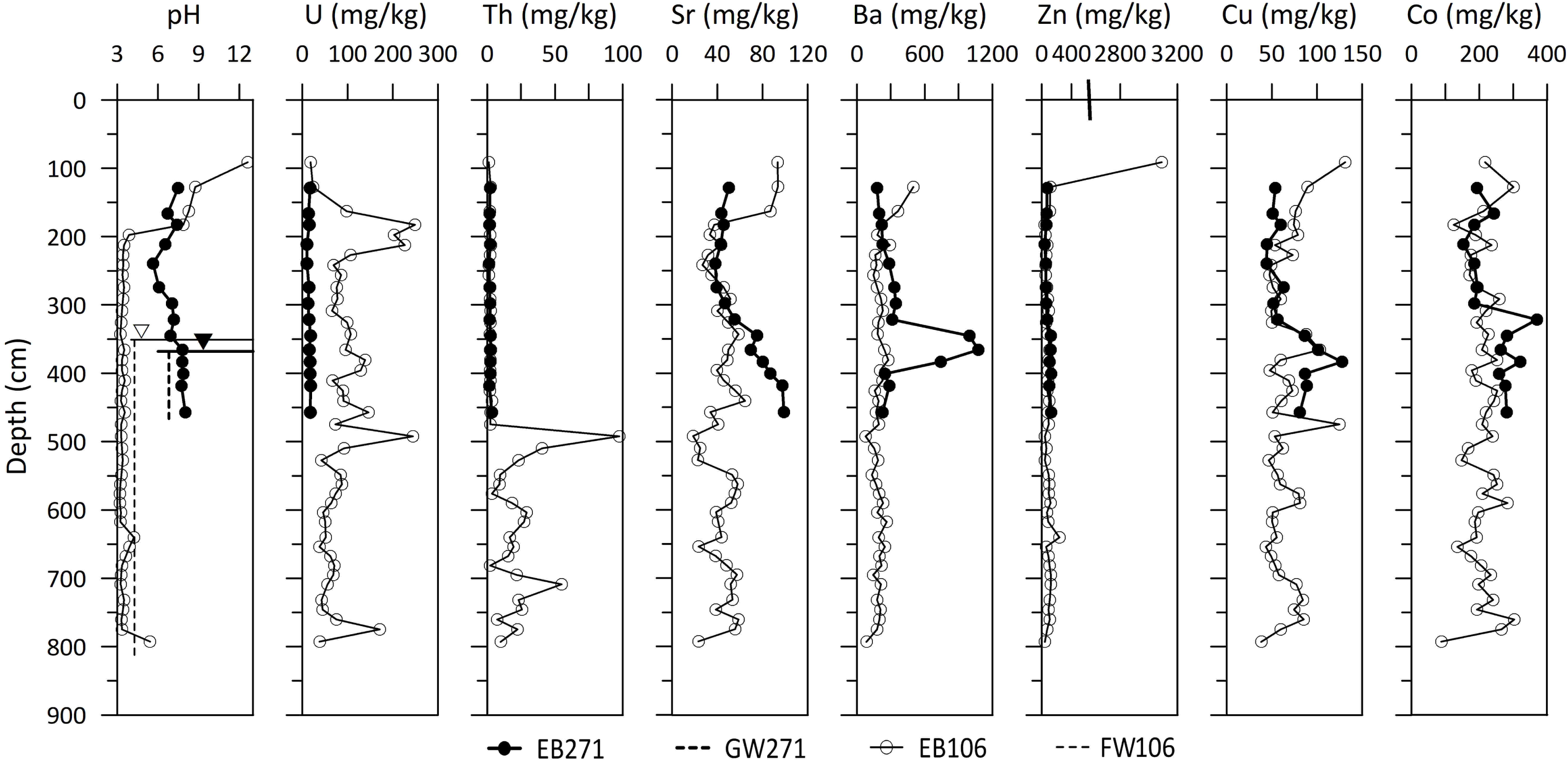
Variation of trace elemental concentrations with depth from core sediments of background EB271 (closed circles and triangles) and contaminated EB106 (open circles and triangles). Open and closed triangles show the water table depth. Vertical broken lines show overall level for groundwater from the nearby well.

Another notable difference between the boreholes is found in the relative levels of Sr and Ba (Fig. 3). The contaminated EB106 had high Sr concentrations (86.9-93.7 mg/kg) in the top three core segments compared to other sections (18.6-64.5 mg/kg) while EB271 exhibited a gradual increase of Sr to 99.1 mg/kg. Ba spiked in EB271 (742-1075 mg/kg) at 345□383 cm depth when approaching the groundwater table, while EB106 showed no such behavior.

Major oxides including SiO_2_, Al_2_O_3_, and Na_2_O did not show obvious trends but CaO, MgO, MnO, and Fe_2_O_3_ were potentially correlated with rainfall events, efforts to control site pH, and changed mineral composition (for example, increases of clay minerals or Fe-Mn oxides) (Fig. 4). The broad MgO peak in contaminated EB106 at 381-441 cm is compatible with the Ba peak of 742□1075 mg/kg at 345-383 cm in that these elements might indicate the increased Ba and Mg rich clay minerals and are likely associated with the water table. K_2_O in EB106 core sediments varied 0.95-5.53 wt.% over 109□334 cm and 1.48□8.03 wt.% over 354-804 cm while at EB271 it was lower at 1.49-3.44 wt.% (Fig. 4). Fe_2_O_3_ and MnO in EB106 had a high correlation with U and Th (Fig. 3 and Fig. 4A). Two high concentration peaks at both 492 cm and 775 cm depth are matched to U and Th peaks. Manganese exhibited a very similar behavior with U rather than iron even under smaller relative content (MnO < 1.5 wt.% and Fe_2_O_3_ < 20 wt.%). This might be due to the closer ionic radius of U (8.7 Å) to Mn (8.1 Å) rather than to Fe (6.9 Å), and the low pH (∼3.5) maintaining the predominantly amorphous Mn phase under pH 4.5 throughout the Eh range (Kim et al., 2009) that can accelerate the similar elemental behavior between U-Mn in contaminated low-pH EB106 core.

**Figure 4.**
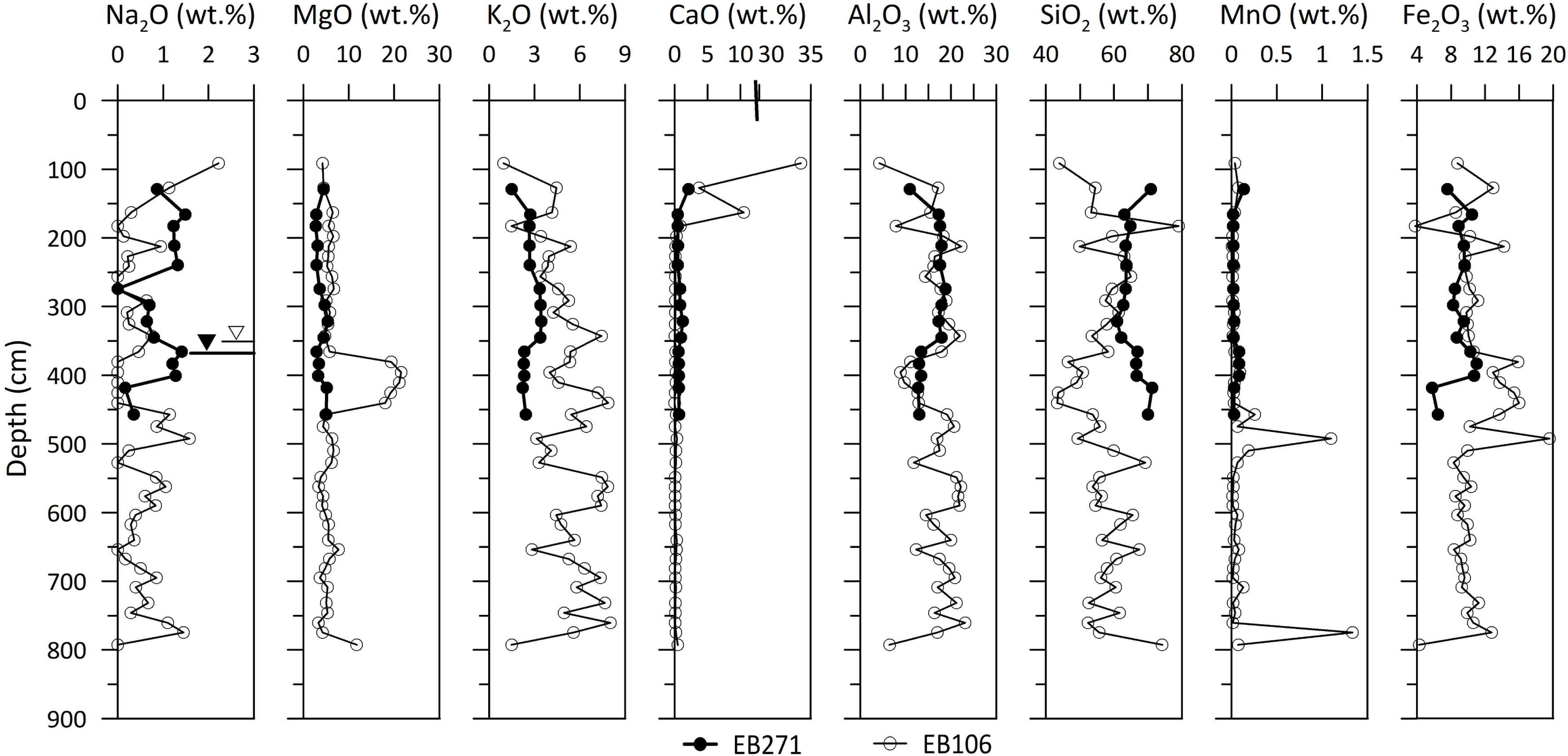
Variation of major oxide component concentrations with depth from core sediments of background EB271 (closed circles and triangles) and contaminated EB106 (open circles and triangles). Open and closed triangles show the water table depth.

The trace and major elemental analysis with ICP-MS following total digestion provided more detailed information of core sediments beyond the total concentration provided by XRF analysis. The total digestion using nitric acid could dissolve more than the surface including short-age ordered mineral phase and amorphous oxides. Even though it could not dissolve completely (i.e., silicate), this fraction includes more indigenous phases. Over 50 metals were analyzed for total metal content by microwave digestion in nitric acidAnalysis of the metal data resulted in several key observations. There were elevated concentrations of several metals including P, Mn, Fe, Co, Ni, As, Cd, and Mo on the interface of the capillary and saturated zones in EB271 between 330 and 350 cm bgs (Fig. 5). These included rare earth, lanthanide and actinides (Ba, La, Pr, Ce, Nd, Sm, Eu, Gd, Dy, Ho, Er, Tm, Yb, Lu; Supp Data #1). In the saturated zone of EB106 at 500 cm bgs there was a similar coinciding elevated metal peak of Be, P, Mn, Fe, Co, U, Pb, Cu, Cr, V, Al, Zn, As, and Mo (Fig. 5; Supp Data #1).

**Figure 5.**
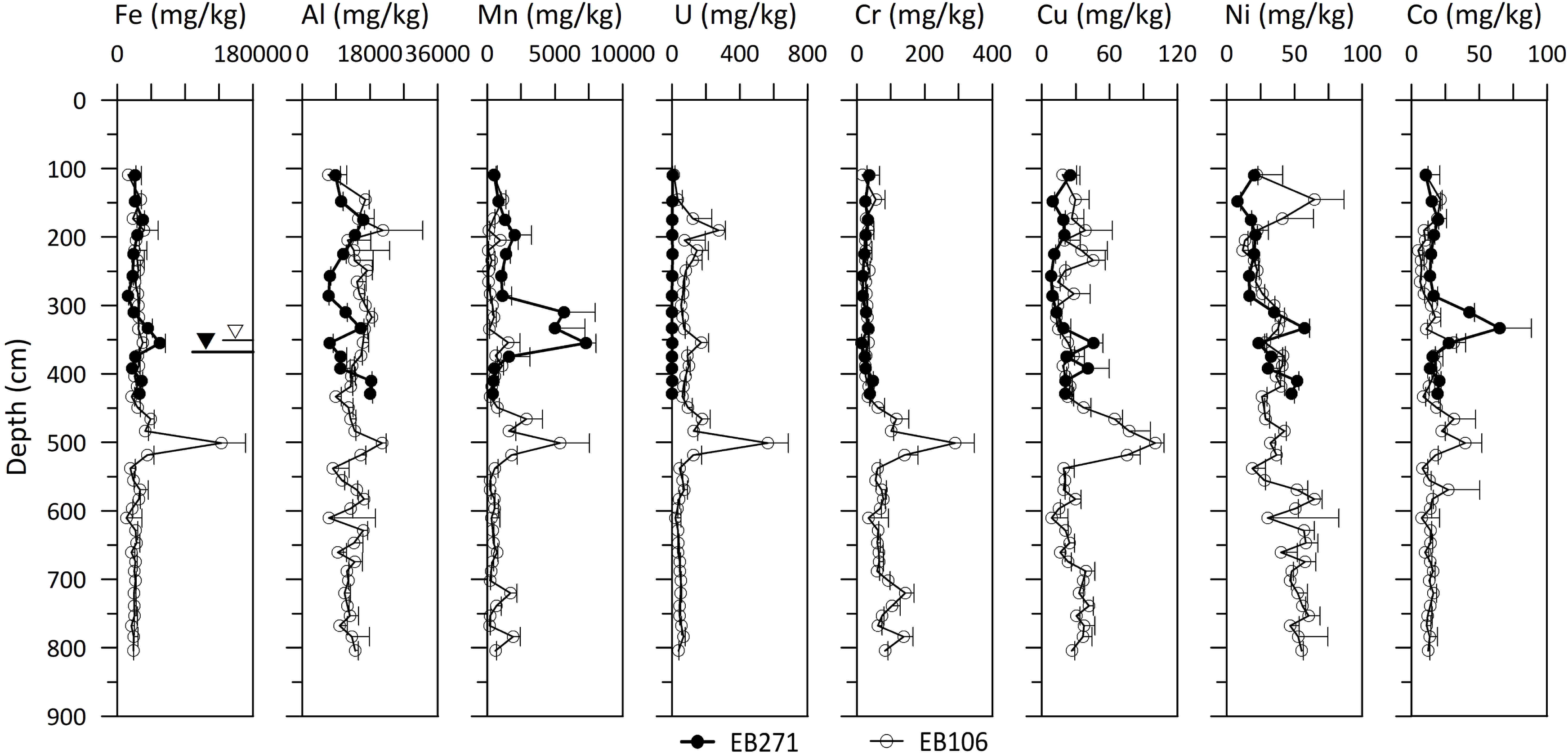
Total metals in sediment segments of EB271 and EB106 by depth. Metals were extracted using microwave digestion of sediment in concentrated nitric acid. Average and standard deviation of three environmental replicates from each homogenized segment are shown. Depth is in centimeters below ground surface (bgs) using inferred values from the middle of each segment.

### 3.2. Inorganic properties of sediment pore water

Extracted sediment pore water ranged from a few microliters to 2 ml (Fig. 6) and this variability was likely due to the high level of physical heterogeneity (Figs. 1 and 2) based on the inherent properties of the sediment and sediment types in the study site (e.g., silty sands versus clays). Extraction volumes were higher above the appearance of the clay mineral-rich layer at ∼400 cm which is less permeable, limiting pore water extraction. This coincided with high contaminant concentrations due to the high surface area of clay minerals and small interstices of sediment particles. Compaction during centrifugation can reduce pore size, limiting water recovery (Jones and Edwards, 1993) so that the extraction amount may not be directly proportional to the total amount of sediment pore water. Hence pore water samples with lower recovered pore water volumes were not analyzed for all parameters due to limited sample and were not plotted in Fig.6. However, the centrifugation method offered a robust, reproducible, and standardized way for determining solute concentration in pore water with quantification of most solutes from organic-poor sediment regardless of centrifuge force and time (Fraters et al., 2017).

**Figure 6.**
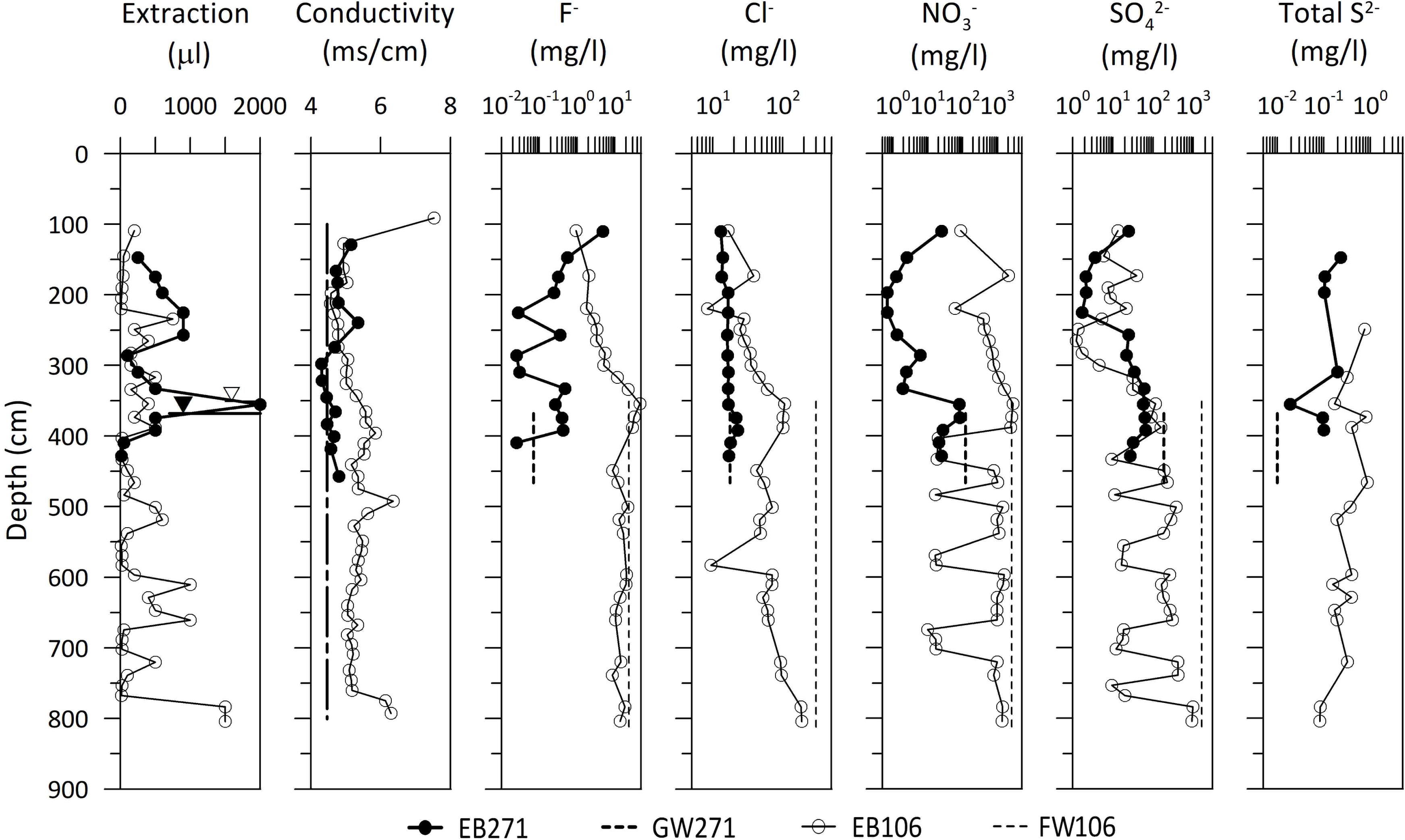
Variation of sediment pore water extraction volume, conductivity, major anions and total sulfide. Conductivity was measured with 10 mM CaCl_2_ buffered solution after pH measurement. Data were plotted with depth from core sediments of background EB271 (closed circles and triangles) and contaminated EB106 (open circles and triangles). Open and closed triangles show the water table depth. Vertical dotted lines represent the properties of groundwater from an adjacent well.

Total conductivity, measured after pH and buffered with 10 mM CaCl_2_ (4.46 mS/cm), exhibited a variation of 4.28-5.35 mS/cm of EB271 and 4.56-6.36 mS/cm except for the top four segments affected by neutralizing process (pH > 7.87). Contaminated EB106 porewater had conductivity above the blank solution due to 2,800 mg/l of nitrate and 980 mg/l of sulfate. The background EB271 were lower than blank concentrations and indicated the potential adsorption of Ca ions onto the background core sediment surface.

Anion concentrations from the contaminated EB106 sediment pore water were considerably higher than from the background EB271 water as expected (Fig. 6). However, EB106 porewater anion levels were lower than in the FW106 groundwater collected. This suggested that groundwater predominantly flowed through preferential paths with high flux and little mixing with water in the interstices of sediment particles, which could impact microbial activity. In contrast, NO_3_^-^ and SO_4_^2-^ closely correlated with the extraction volume, especially below the groundwater table at 501-538 cm, 597-661 cm, and 720-739 cm bgs. This also indirectly suggests high groundwater flux with preferential flow derived from different subsurface geological media. Nitrate remained high throughout in EB106 pore water, while sulfate concentration was low (1.2-127 mg/l) in the vadose zone samples compared to the groundwater (1,630 mg/l). This suggests that sulfate-reducing bacteria may be more active in areas with detectable sulfide (< 1 mg/l), similar to EB271 but not for EB106.

### 3.3. Organic acid in sediment pore water in core sediments

Acetate is a key intermediate produced during the turnover of the sediment organic carbon under the well-drained and short-term anoxic conditions (Küsel et al., 2002). Formate is generally regarded as a waste product of fermentation (i.e. *Escherichia coli*) (McDowall et al., 2014). Sediment pore water acetate and formate concentrations both decreased at the water table in EB271 while no variation was noted for EB106. The latter showed a steady decrease in the saturated zone (Fig. 7).

**Figure 7.**
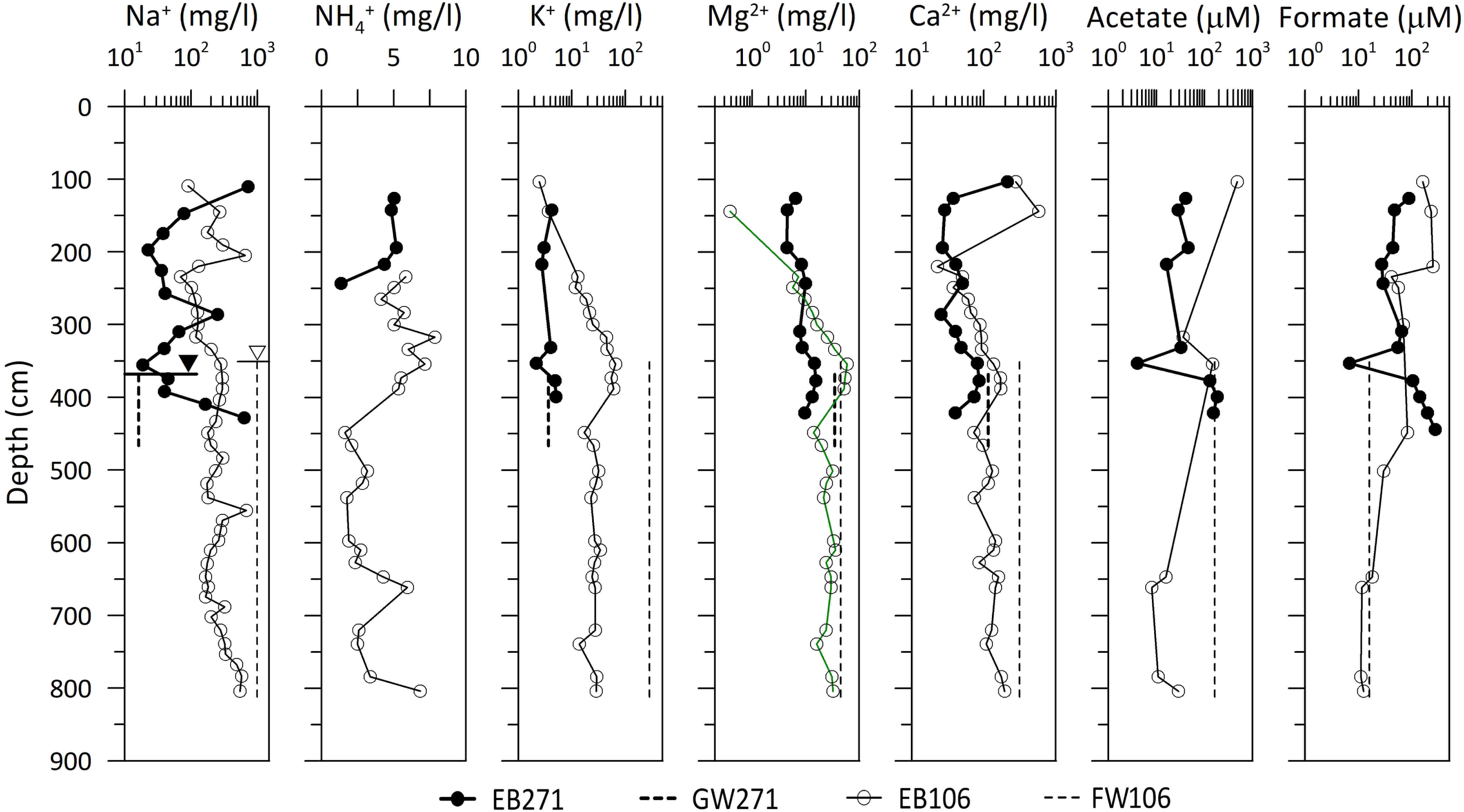
Variation of major cations and organic acid extracted sediment pore water. Data were plotted with depth from core sediments of background EB271 (closed circles and triangles) and contaminated EB106 (open circles and triangle). Open and closed triangles show the water table depth. Vertical dotted lines represented the properties of contaminated groundwater.

### 3.4. Microbial signature in core sediments

As shown in Fig. 8, several variations appeared when comparing direct cell counts using AODC to core sediment (counts/g) compared to groundwater (counts/ml). These included higher numbers in the vadose zone of EB106, consistent with previous studies (Christensen et al., 2018), and is likely due to the presence of oxygen, a neutral pH and lower contaminant levels.

**Figure 8.**
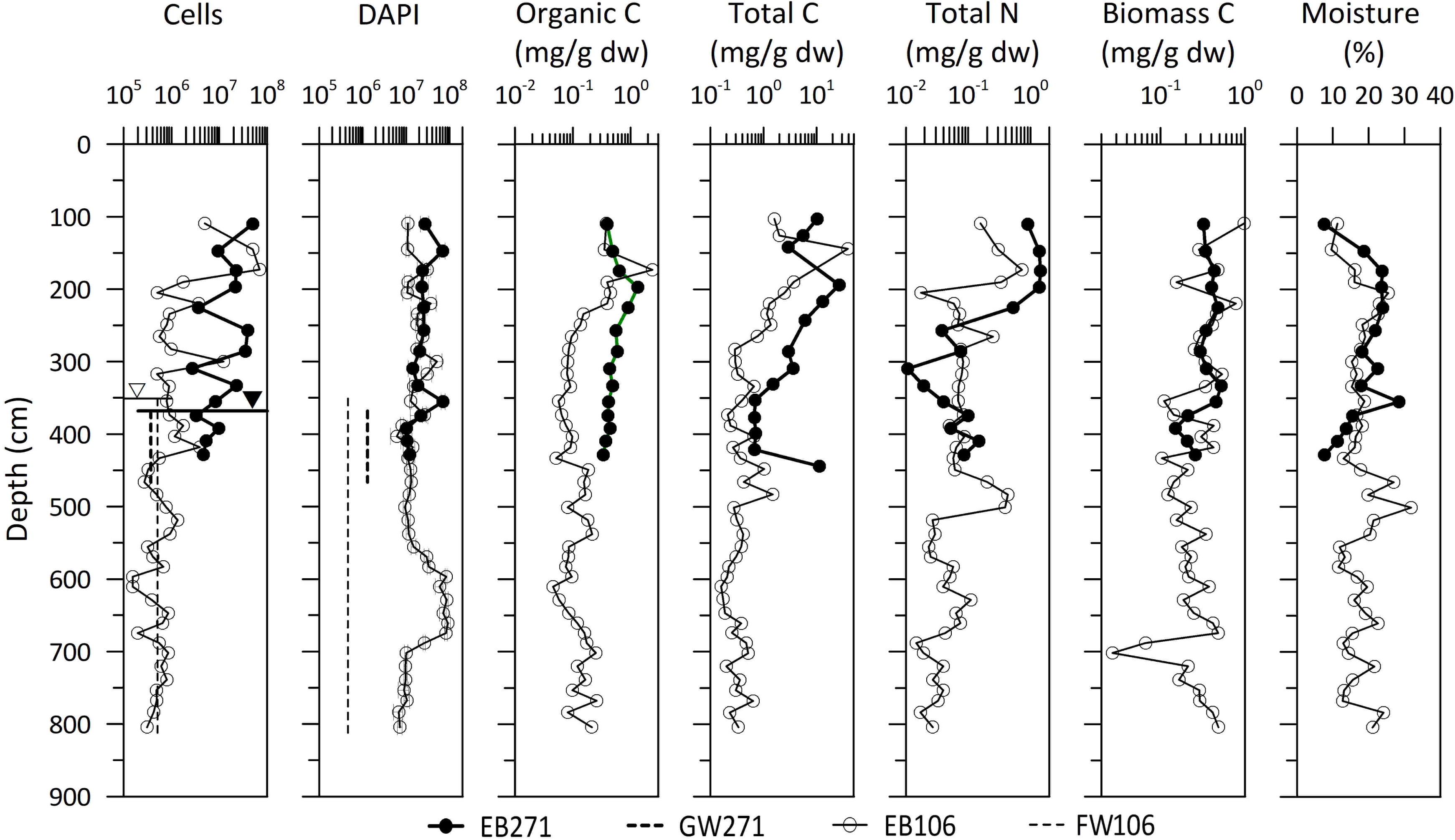
Variation of direct cell counting, organic carbon, total carbon, total nitrogen, biomass carbon using a chloroform fumigation extraction, and sediment moisture. Data were plotted with depth from core sediments of background EB271 (closed circles and triangles) and contaminated EB106 (open circles and triangles). Open and closed triangles show the water table depth. Vertical dotted lines represented the properties of contaminated groundwater. Cell numbers are in counts/g and counts/ml for sediment and groundwater, respectively.

### 3.5. Sediment gas

Selected core sediments along the depth selected from vadose, capillary, and saturated zone were amended with groundwater from neighboring monitoring wells (GW271 and FW106) and exhibited two distinct developments in the gas phases (Fig. 9). Background EB271 sediments showed high biomass (Fig. 8) and generated more CH_4_ at up to 0.56 mM/day/g compared to EB106 (up to 0.02 mM/day/g). In contrast, EB106 produced consistent and higher CO_2_ concentrations (up to 19 mM/day/g) compared to 3.2 mM/day/g in EB271. Higher nitrous oxide was detected in EB106, consistent with higher nitrate (Fig. 6). EB271 core samples had only one sample above detection out of six selected samples.

**Figure 9.**
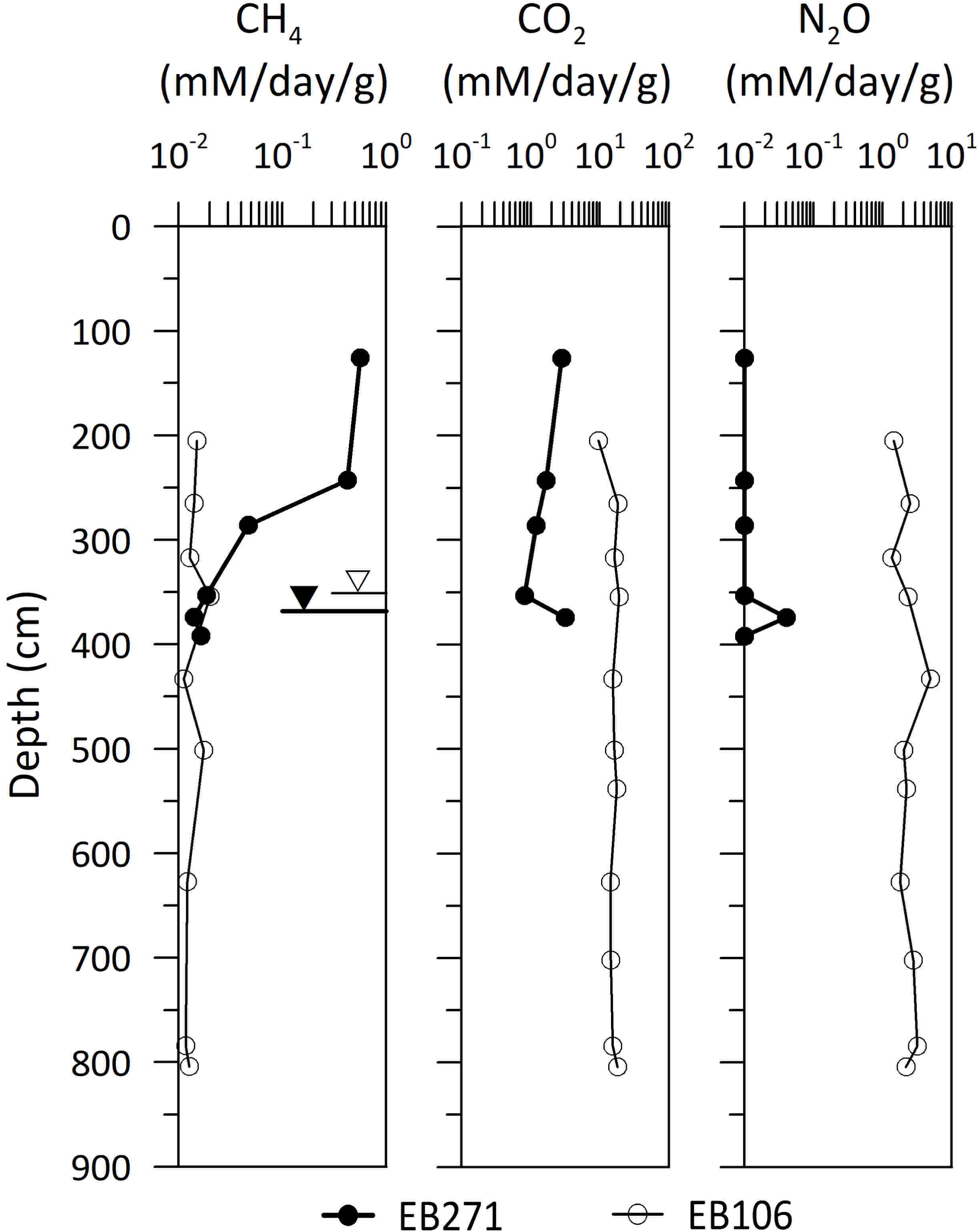
Variation of biogenic CH_4_, CO_2_, and N_2_O from the anaerobic incubations of core sediment from vadose, capillary and saturated zones, respectively, and unfiltered groundwater at room temperature in the dark for 4 weeks. Data were plotted with depth from core sediments of background EB271 (closed circles and triangles) and contaminated EB106 (open circles and triangles). Open and closed triangles show the water table depth.

### 3.6. Sediment mineralogy

Sediment minerals were identified and semi-quantified based on the XRD patterns from the selected sediment samples (Supplementary Fig. 1). To compare with other results, the samples as for gas analysis were used, however some high U-concentration samples were omitted due to lab safety restrictions. Major mineral constituents were common sediment minerals of quartz, Na-feldspar (NaAlSi_3_O_8_), K-feldspar (KAlSi_3_O_8_), illite (muscovite), and kaolinite (Fig. 10). Only top sediment of EB106 which had high pH (∼13) from surface neutralization included calcite (CaCO_3_). Without any further process in terms of oxalate treatment for amorphous phase and citrate-bicarbonate-dithionate treatment for crystalline iron oxide (Kim et al., 2009), dominant peaks from accessory minerals were not identifiable.

**Figure 10.**
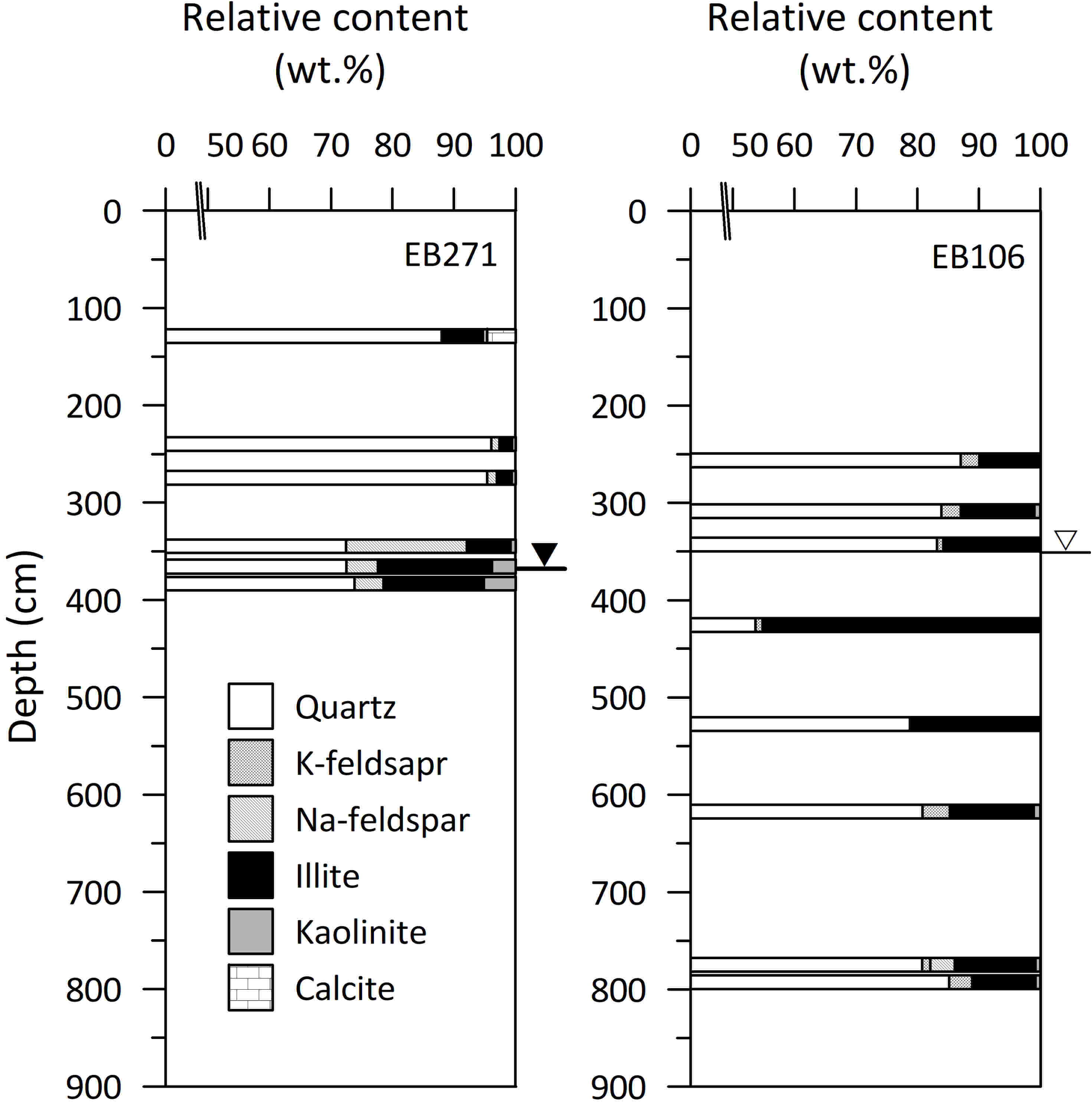
Variation of relative content of mineral constituents from the XRD analysis on selected samples among gas analysis.

Semi-quantification of bulk sediment showed that the vadose zone of EB271 with high biomass was high in quartz (88-96 wt.%) with clay minerals (illite + kaolinite, 2.6-7.4 wt.%). High concentrations of trace elements (U, Th, Fe, and Mn) are closely related to clay-rich layer (quartz, 54-79 wt.% with clay minerals, 21□45 wt.%) in EB106. High concentrations U or Th were highly correlated to clay minerals or near bedrock which prevented the vertical movement of water (quartz, 73-74 wt.% with clay minerals, ∼22 wt.%).

## 4. DISCUSSION

### 4.1. Sediment characterization

The variability of moisture below the water table Fig. 2, may be partially attributed to sediment heterogeneity since gravels and sands can contain more free water than lower specific yield sediments such as silts and clays (Fetter, 2001). Except for the first meter, the predominant mean grain size in both boreholes was generally > 2.5 mm (i.e., coarse sand). The *in situ* visual and manual classification of sediments from both boreholes (Fig. 1 and Fig. 2) indicated a high level of physical heterogeneity and suggested that specific yield and subsequent groundwater flow are highly variable with respect to depth within each borehole. Highly variable groundwater flow, with respect to depth, is a common occurrence at sites with a high level of physical heterogeneity and a wide variety of down-well devices are available to quantitatively document its occurrence (Bayless et al., 2011).

The broad U peak in EB106 at 183-212 cm without a corresponding Th peak was likely caused by the sediment pH; the pH ranged from 5.62-8.01 along EB271 and from 3.18-12.62 along EB106 (Fig. 3). Except for the four top core segments with high pH (7.87-12.62), the remainder of EB106 sediment fell in a pH range of 3.18-5.40. The high pH values of EB106 at the top core segments might be due to the application of calcareous materials used in attempts at pH neutralization. This is in agreement with results from a previous study (Ahmed et al., 2014) showing that Th solubility was 5-14 times greater than uranium in acidic sediments (pH 3.6-4.7). In contrast, U was significantly more soluble than Th in mildly acidic sediments (pH 5.8-7.0). Hence the enhanced solubility of Th at this low pH range before the neutralization facilitates Th leaching and only a broad U peak observed at 183-212 cm bgs. The solubility and mobility of U and Th can also be influenced by organic matter (Ahmed et al., 2014), however previous research at the ENIGMA field research site indicated that the sediment is usually limited for electron donors and organic nutrients (Gu et al., 2003). Therefore, pH is likely the main reason for the behavior of U and Th.

High Sr and Ba concentrations were also noted in the upgradient sediment of EB271, suggesting the high Sr and Ba are more likely related to the interaction between soil minerals close to groundwater in the interstices of the soil particles. The relatively lower concentrations of Sr and Ba in the lower levels of EB106 sediment are likely a direct effect of the pH. An abnormally high pH near the surface of EB106 implies that the surface reclamation with limestone neutralized the acidic contaminated area since Sr is a dominant replaceable element to Ca in carbonate minerals due to their similar charge and ionic radius (Blundy and Wood, 1991). Among the transitional metals, Zn was largely absent in both cores while Cu and Co (Fig. 3) exhibited a weak but variable increase near the water table and close to bedrock. These trends agree with the contaminants enriching at the soil-water interfaces (Milligan and Law, 2013) or water-air interfaces (Zhang and Christopher, 2016). CaO levels reflected the neutralizer application at the surface of EB106 and was quickly lost in the study site shales, siltstone, and limestone (Hatcher et al., 1992) due to the low pH. The higher levels of K_2_O in EB106 may be from acidic chemical weathering of saprolite, resulting in leached clay minerals. This is in agreement with the previous study finding that illite was a dominant clay mineral species and microcline (potassium feldspar) under the water table in the weathered saprolite zone (Moon et al., 2006). U was found to be highly adsorbed or incorporated into amorphous manganese oxide (Moon et al., 2006), likely sourced from Mn-rich muscovite in the shale and Mn-rich biotite in the blackish band of the limestone (Kim et al., 2009). In this site, clay minerals are heavily coated with Fe- and Mn-oxides as well as most of the fracture pathways and matrix blocks are coated with amorphous Fe- and Mn-oxides. The redox environment and the presence of redox reactive minerals and solids (e.g. Mn-oxides and organic matter) also impact contaminant transport at this site (Watson et al., 2004).

As shown in Fig. 5, Mn Co, and Ni from EB271 background borehole had peaks matching well with the groundwater table. In contrast, many metals including Be, P, Mn, Fe, Co, U, Pb, Cu, Cr, V, Al, Zn, As, and Mo (Fig. 5; Supp Data #1) had peaks well below the groundwater table, especially at 500 cm bgs, in the anthropogenically contaminated EB106 in the borehole. The segment at 500 cm bgs also had decreased density and increased moisture and conductivity (Figure 2 and 6) as well as visible metal bearing particles shown in Figure 11. These findings suggest an increase contaminated groundwater flow from the S-3 ponds at this depth, supporting the assertion of preferential flow paths in EB106 area.

**Figure 11.**
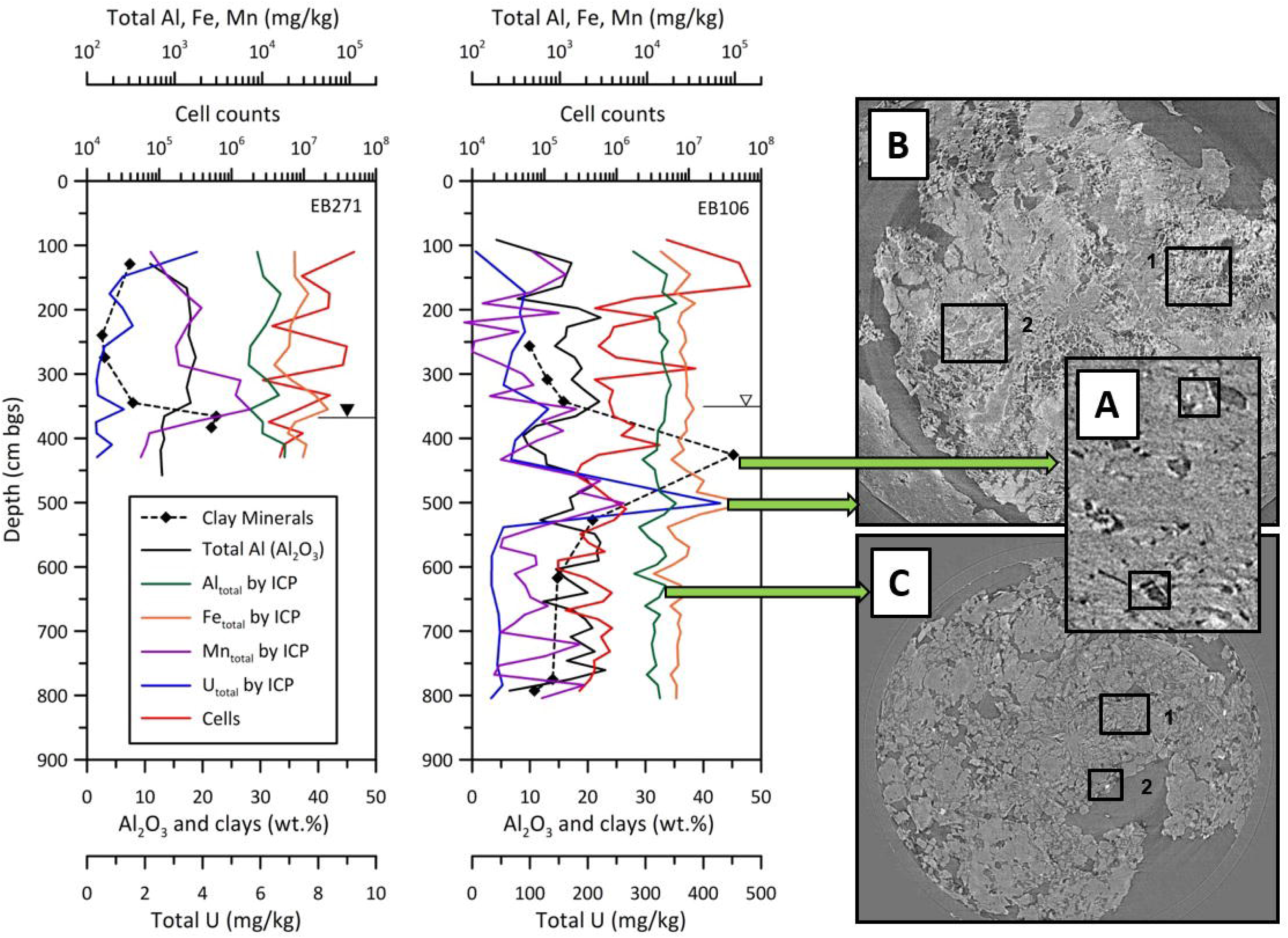
Correlation between uranium and the relative content of clay minerals (illite and kaolinite) among mineral constituents, total Al, Fe and Mn concentration, total aluminum content, and cell counts. Micro-computerized tomographic images provide direct observation of solid geochemical condition; A (EB106 section 5-5): spine structure of clay (illite); B (EB106 section 6-3): 1, low porosity section with webby structure of clay (smectite), 2, abundant white seam of metals; C (EB106 section 7-6): 1, bedrock fragment, 2 metal bearing particles.

### 4.2. Inorganic properties of sediment pore water

The most dominant difference among the major cation measurements is the occurrence of NH_4_^+^ in the sediment pore water that was not detected in the collected groundwater from either of the wells located next to the boreholes (Fig 7). This suggests that the porewater is reflective of anaerobic niches in the sediment while the bulk groundwater environment is aerobic or microaerophilic. Potassium and Mg^2+^ in EB106 exhibited a broad peak at the groundwater table. As shown in the conductivity profiles (Fig. 5), background EB271 had lower conductivity ranges than the 10 mM CaCl_2_ blanks indicating the loss of ions by adsorption and absorption. However, contaminated EB106 showed higher conductivity with a sharp peak matching U, Th, Fe, and Mn peaks. Therefore, increased K^+^ and Mg^2+^ might be released by weathering of clay minerals. Kim et al. interpreted the elemental cycles of K leaching by weathering pathways (muscovite from shale → illite → kaolinite) and that of Fe and Mg (biotite and chlorite from interbedded limestone → vermiculite → kaolinite) (Kim et al., 2009). The enriched alkali and alkaline earth elements in the sediment pore water likely reflected the increased trace elements as contaminants around the groundwater table produced by cation exchange with transitional metals with low solubility products which could produce more soluble oxide or amorphous phases, although the whole core sections had a low pH of ∼3.5. Distinctively high Ca^2+^ might be related to the artificial neutralization (limestone, soil, and saprolite) from construction activity (Moon et al., 2006), however the behavior of Na^+^ is not certain. The peaks of 205 cm and 556 cm from EB106 and 286 cm from EB271 did not match any peaks among the analyzed items.

### 4.3. Organic acid in sediment pore water in core sediments

Acetate was also lower in depth specific measurements than the overall groundwater, implying again that microenvironments within sediment particles may be preventing direct exposure to toxic contaminants while allowing for the consumption of acetate, possibly combined with sulfur respiration (Pfennig and Biebl, 1976). This emphasizes that the sample for isolation or interpretation likely can be misinterpreted by the dissolved chemical species and organic carbon balances in the groundwater versus depth dependent measurements.

### 4.4. Microbial signature in core sediments

Cell counts from both core sediments had higher populations than from groundwater, presumably since biofilms on the sediment surface might account for a large fraction of microbial populations. Higher cell counts in EB271 overall are also likely due to less harsh conditions, which likely reflects a microbial community with more genomic functional redundancy than the stressed community (Hemme et al., 2015), however these harsh conditions may have provided for specific niches for microbial communities with unique metabolic capabilities (Smith et al., 2012). Overall, all carbon and nitrogen and moisture levels were similar between the cores.

### 4.5. Sediment gas

The higher nitrous oxide in EB106 may be partially due to abiotic denitrification at low pH in the presence of Fe. This release does not correlate with either cell counts or concentrations of oxidized nitrogen species and might thus be limited to the mechanism of abiotic chemodenitrification. The peak of nitrous oxide release found below the water table in EB271 sediment coincides with high levels of oxidized nitrogen species found in sediment pore water.

### 4.6. Sediment mineralogy

The combined results among the clay minerals, water, and trace element contaminants are also confirmed by microCT images. As shown in Fig. S2a, the brighter portion of images of EB106 6-3 core section showed the presence of heavy metals like U, Th, Fe, and Mn that matched with the abnormal peaks from the XRF EB106 6-3 sample at 492□510 cm bgs (Figs. 3 and 4). These images also exhibited a honeycomb structure suggesting the presence of clay minerals. XRD analysis was not performed on section 6-3 samples due to high U concentrations. The most enriched clay minerals were found in section EB106 5-5 by XRD (Fig. 11). MicroCT images from this section (Fig. S2b) for slices #1907, #1942, and #1979 exhibited clay mineral features. MicroCT images were also obtained for the next clay enriched section analyzed by XRD, 6-5 (Fig. S2c).

These findings emphasize the notion that limited chemical analyses of major or trace elements (i.e. XRF analysis) cannot fully determine the reasons for contaminant enrichment at a specific depth. For example, Al_2_O_3_ showed the consistent weight fraction and could not be separated from feldspars to obtain the clay amount. Direct mineralogical analysis followed by semi-quantification revealed the clay-rich layer occupies a large surface area, has less permeability, and a higher adsorption of contaminants. Detectable calcite in EB106 was compatible with the high Sr concentration among trace elements. Also, most minerals are end-members, therefore high concentration of Ca, Mg, Ba, and Sr could be in present in calcite and other carbonate minerals (i.e. norsethite, [MgBa(CO_3_)_2_], even though the amount was below XRD detection.

## 5. CONCLUSIONS

The present study compared fresh sediment cores from upgradient and contaminated areas with similar characteristics. When exploring microbial communities in the subsurface, consideration of a wide spectrum of unique sample points is required rather than simply the averaged and homogenized groundwater chemistry. Consideration of geochemical parameters as a function of depth can reveal complicated but meaningful relationships between geochemical and hydrological conditions that may well influence microbial community structure and function. Groundwater analysis of nearby wells only revealed high sulfate and nitrate concentrations while the same analysis using sediment pore water samples with depth was able to pinpoint areas likely high in sulfate- and nitrate- reducing bacteria based on their decreased concentration and production of reduced by-products that could not be seen in the groundwater samples. This emphasizes that the sample for isolation or interpretation likely can be misinterpreted by the dissolved chemical species and organic carbon balances in the groundwater versus depth dependent sediment measurements.

Low flow (i.e. dry season) at the time of sampling provided the maximum time for a distinct community to develop on particles (Enzien et al., 1994; LaMontagne and Holden, 2003). However, metabolic rates were found to be higher in a shallow sandy aquifer compared with a confined clayey aquifer likely due to the reduced interconnectivity followed by a reduction in microbial and nutrient mobility (Chapelle and Lovley, 1990). Therefore, geochemical data from sediment pore water and solid media should be merged with mineralogical findings. Positive correlations among pore water content, total organic carbon, trace metals and clay minerals were previously observed (Böttcher et al., 2000) and the present study revealed a more complicated relationship; among contaminant, sediment texture, groundwater table, and biomass and the need for more microbial analysis of these samples. A merged mineralogical investigation with semi-quantification could reveal the abundant clay minerals with high surface area and high water-holding capacity of micro-pores of the fine clay rich layer under the interference of feldspars. Such geological features can impede the movement of nutrients from the surface.

At the groundwater table of EB106, trends followed the previous study (Chapelle and Lovley, 1990) with cell counts decreasing while clay content and Al_2_O_3_ increased and resulted in low porewater extraction. This fluctuating capillary interface maintained high concentration of Fe and Mn-oxides combined with trace elements including U, Th, Sr, Ba, Cu, and Co. This indicates that the mobility of highly toxic elements, sediment structure, and biogeochemical factors are all linked together in their impact on microbial communities, emphasizing that solid interfaces play an important role in determining the abundance of bacteria in the sediments (Lovley, 1998; Paradis et al., 2018). In contrast, the background EB271 had similar Al_2_O_3_ content over the entire core. While this decreased below 400 cm, Al_2_O_3_ detection was impaired by the presence feldspars, and therefore only mineralogical analysis revealed the clay mineral’s important role with sediment texture. This suggested that groundwater predominantly flowed through preferential paths with high flux and little mixing with water in the interstices of sediment particles, which could impact microbial activity.

## ACKNOWLEDGEMENTS

We thank Kelley Meinhardt in the Stahl lab for performing the sediment gas measurements and Deanne Brice at ORNL for C&N analysis. This material by ENIGMA-Ecosystems and Networks Integrated with Genes and Molecular Assemblies (http://enigma.lbl.gov), a Scientific Focus Area Program at Lawrence Berkeley National Laboratory is based upon work supported by the U.S. Department of Energy, Office of Science, Office of Biological & Environmental Research under contract number DE-AC02-05CH11231. Oak Ridge National Laboratory is managed by UT-Battelle, LLC, for the U.S. Department of Energy under contract DE-AC05-00OR22725.

## References

Ahmed, H., Young, S.D. and Shaw, G. (2014) Factors affecting uranium and thorium fractionation and profile distribution in contrasting arable and woodland soils. Journal of Geochemical Exploration 145, 98–105.

Ahn, J.H. (1992) Application of an-XRD-pattern calculation method to quantitative analysis of clay minerals. Journal of The Minieralogical Society of Korea 5, 32–41.

Bayless, E.R., Mandell, W.A. and J.R., U. (2011) Accuracy of flowmeters measuring horizontal groundwater flow in an unconsolidated aquifer simulator. Groundwater Monitoring and Remediation 31, 48–62.

Blundy, J.D. and Wood, B.J. (1991) Crystal-chemical controls on the partitioning of Sr and Ba between plagioclase feldspar, silicate melts, and hydrothermal solutions. Geochimica Cosmochimica Acta 55, 193–209.

Böttcher, M.E., Hespenheide, B., Llobet-Brossa, E., Beardsley, C., Larsen, O., Schramm, A., Wieland, A., Böttcher, G., Berninger, U. and Amann, R. (2000) The biogeochemistry, stable isotope geochemistry, and microbial community structure of a temperate intertidal mudflat: an integrated study. Continental Shelf Research 20, 1749–1769.

Chapelle, F.H. and Lovley, D.R. (1990) Rates of microbial metabolism in deep coastal plain aquifers. Applied and Environment Microbiology 56, 1865–1875.

Christensen, G.A., Moon, J., Veach, A.M., Mosher, J.J., Wymore, A.M., van Nostrand, J.D., Zhou, J.Z., Hazen, T.C., Arkin, A.P. and Elias, D.A. (2018) Use of in-field bioreactors demonstrate groundwater filtration influences planktonic bacterial community assembly, but not biofilm composition. Plos One 13, 20.

Chung, F.H. (1974) Qunatitative Interpretation of X-ray diffraction patterns of mixtures. II. Adiabatic principle of X-ray diffraction analysis of mixtures. Journal of Applied Crystallography 7, 526–531.

De Goffau, A., Van Leeuwen, T.C., Van den Ham, A., Doornewaard, G.J. and Fraters, B. (2012) Minerals Policy Monitoring Programme Report 2007-2010, Methods and Procedures. National Institute for Public Health and the Environment.

Enzien, M.V., Picardal, F., Hazen, T.C., Arnold, R.G. and Fliermans, C.B. (1994) Reductive Dechlorination of Trichloroethylene and Tetrachloroethylene under Aerobic Conditions in a Sediment Column. Applied and Environmental Microbiology 60, 2200–2204.

EPA, U.S. (2007) Method 3051A (SW-846): Microwave assisted acid digestion of sediments, sludges, and oils, Revision 1 ed. U.S. EPA, Washington, D.C.

Fetter, C.W. (2001) Applied Hydrogeology, 4th ed. Prentice Hall, Upper Saddle River, NJ.

Fraters, D., Boom, G.J.F.L.B. L.J.M., de Weerd, H. and Wolters, M. (2017) Extraction of soil solution by drainage centrifugation—effects of centrifugal force and time of centrifugation on soil moisture recovery and solute concentration in soil moisture of loess subsoils. Environmental Monitoring and Assessment 189, 83.

Ge, X.X., Vaccaro, B.J., Thorgersen, M.P., Poole, F.L., Majumder, E.L., Zane, G.M., De Leon, K.B., Lancaster, W.A., Moon, J.W., Paradis, C.J., von Netzer, F., Stahl, D.A., Adams, P.D., Arkin, A.P., Wall, J.D., Hazen, T.C. and Adams, M.W.W. (2019) Iron- and aluminium-induced depletion of molybdenum in acidic environments impedes the nitrogen cycle. Environmental Microbiology 21, 152–163.

Griebler, C. and Lueders, T. (2009) Microbial biodiversity in groundwater ecosystems. Freshwater Biology 54, 649–677.

Gu, B., Brooks, S.C., Roh, Y. and Jardine, P.M. (2003) Geochemical reactions and dynamics during titration of a contaminated groundwater with high uranium, aluminum, and calcium. Geochimica Cosmochimica Acta 67, 2749–2761.

Hatcher, R.D., Lemiszki, P.J., Dreier, R.B., Ketelle, R.H., Lee, R.R., Leitzke, D.A., McMaster, W.M., Foreman, J.L. and Lee, S.Y. (1992) Status report on the geology of the Oak Ridge Reservation.

Hazen, T.C., Jimenez, L., Devictoria, G.L. and Fliermans, C.B. (1991) Comparison of Bacteria from Deep Subsurface Sediment and Adjacent Groundwater. Microbial Ecology 22, 293–304.

Hemme, C.L., Deng, Y., Gentry, T.J., Fields, M.W., Wu, L., Barua, S., Barry, K., Tringe, S.G., Watson, D.B., He, Z., Hazen, T.C., Tiedje, J.M., Rubin, E.M. and Zhou, J. (2010) Metagenomic insights into evolution of a heavy metal-contaminated groundwater microbial community. The ISME Journal 4, 660.

Hemme, C.L., Tu, Q., Shi, Z., Qin, Y., Gao, W., Deng, Y., Van Nostrand, J.D., Wu, L., He, Z., Chain, P.S.G., Tringe, S.G., Fields, M.F., Rubin, E.M., Tiedje, J.M., Hazen, T.C., Arkin, A.P. and Zhou, J. (2015) Comparative metagenomics reveals impact of contaminants on groundwater microbiomes. Frontiers in Microbiology 6, 1205.

Jones, D.L. and Edwards, A.C. (1993) Effect of moisture content and preparation technique on the composition of soil solution obtained by centrifugation. Communications in Soil Science and Plant Analysis 24, 177–186.

Kim, Y.-J., Moon, J.-W., Roh, Y. and Brooks, S.C. (2009) Mineralogical characterization of saprolite at the FRC background site in Oak Ridge, Tennessee. Environmental Geology 58, 1301–1307.

Küsel, K., Wagner, C., Trinkwalter, T., Gößner, A.S., Bäumler, R. and Drake, H.L. (2002) Microbial reduction of Fe(III) and turnover of acetate in Hawaiian soils. FEMS Microbiology Ecology 40, 73–81.

LaMontagne, M.G. and Holden, P.A. (2003) Comparison of free-living and particle-associated bacterial communities in a coastal lagoon. Microbial Ecology 46, 228–237.

Lovley, D.R. (1998) Geomicrobiology: Interactions between microbes and minerals. Science 280, 54–55.

McDowall, J.S., Murphy, B.J., Haumann, M., Palmer, T., Armstrong, F.A. and Sargent, F. (2014) Bacterial formate hydrogenlyase complex. Proceedings of the National Academy of Sciences, USA 111, 3948–3956.

Milligan, T.G. and Law, B.A. (2013) Contaminants at the sediment-water interface: Implications for environmental impact assessment and effects monitoring. Environmental Science & Technology 47, 5828–5834.

Moon, J.-W., Ivanov, I.N., Joshi, P.C., Armstrong, B.L., Wang, W., Jung, H., Rondinone, A.J., Jellison Jr., G., Meyer III, H.M., Jang, G.G., Meisner, R.A., Duty, C.E. and Phelps, T.J. (2014) Scalable production of microbially-mediated ZnS nanoparticles and application to functional thin films. Acta Biomaterialia 10, 4474–4483.

Moon, J.-W., Roh, Y., Phelps, T.J., Phillips, D.H., Watson, D.B., Kim, Y.-J. and Brooks, S.C. (2006) Physicochemical and mineralogical characterization of soil–saprolite cores from a field research site, Tennessee. Journal of Environmental Quality 35, 1731–1741.

Moon, J.-W., Song, Y., Moon, H.-S. and Lee, G.H. (2000) Clay minerals from tidal flat sediments at Youngjong Island, Korea, as a potential indicator of sea-level change. Clay Minerals 35, 841–855.

Paradis, C.J., McKay, L.D., Perfect, E., Istok, J.D. and Hazen, T.C. (2018) Push-pull tests for estimating effective porosity: expanded analytical solution and in situ application. Hydrogeology Journal 26, 381–393.

Pfennig, N. and Biebl, H. (1976) *Desulfuromonas acetoxidans* gen. nov. and sp. nov., a new anaerobic, sulfur-reducing, acetate-oxidizing bacterium. Archives of Microbiology 110, 3–12.

Sinclair, J.L. and Ghiorse, W.C. (1989) DISTRIBUTION OF AEROBIC-BACTERIA, PROTOZOA, ALGAE, AND FUNGI IN DEEP SUBSURFACE SEDIMENTS. Geomicrobiology Journal 7, 15–31.

Smith, H.J., Zelaya, A.J., De Leon, K.B., Chakraborty, R., Elias, D.A., Hazen, T.C., Arkin, A.P., Cunningham, A.B. and Fields, M.W. (2018) Impact of hydrologic boundaries on microbial planktonic and biofilm communities in shallow terrestrial subsurface environments. Fems Microbiology Ecology 94, 16.

Smith, M.B., Rocha, A.M., Chris S. Smillie, C.S., Olesen, S.W., Paradis, C., Wu, L., Campbell, J.H., Fortney, J.L., Mehlhorn, T.L., Lowe, K.A., Earles, J.E., Phillips, J., Techtmann, S.M., Joyner, D.C., Elias, D.A., Bailey, K.L., Hurt, J. R.A., Preheim, S.P., Sanders, M.C., Yang, J., Mueller, M.A., Brooks, S.C., Watson, D.B., Zhang, P., He, Z., Dubinsky, E.A., Adams, P.D., Arkin, A.P., Fields, M.W., Zhou, J., Alm, E.J. and Hazen, T.C. (2015) Natural bacterial communities serve as quantitative geochemical biosensors. mBio 6, e00326–00315.

Smith, R.J., Jeffries, T.C., Roudnew, B., Fitch, A.J., Seymour, J.R., Delpin, M.W., Newton, K., Brown, M.H. and Mitchell, J.G. (2012) Metagenomic comparison of microbial communities inhabiting confined and unconfined aquifer ecosystems. Environmental Microbiology 14, 240–253.

Watson, D.B., Kostka, J.E., Fields, M.W. and Jardine, P.M. (2004) The Oak Ridge Field Research Center Conceptual Model. Oak Ridge National Laboratory, Oak Ridge, p. 55.

Zhang, Z. and Christopher, G. (2016) Effect of particulate contaminants on the development of biofilms at air/water interfaces. Langmuir 32, 2724–2730.

