## Supplemental Table 1-4 for "Integrated characterization of subsurface media from locations up- and down-gradient of a uranium-contaminated aquifer"

*This manuscript has been authored by UT-Battelle, LLC under Contract No. DE-AC05-00OR22725 with the U.S. Department of Energy. The United States Government retains and the publisher, by accepting the article for publication, acknowledges that the United States Government retains a non-exclusive, paid-up, irrevocable, world-wide license to publish or reproduce the published form of this manuscript, or allow others to do so, for United States Government purposes. The Department of Energy will provide public access to these results of federally sponsored research in accordance with the DOE Public Access Plan (http://energy.gov/downloads/doe-public-access-plan).*

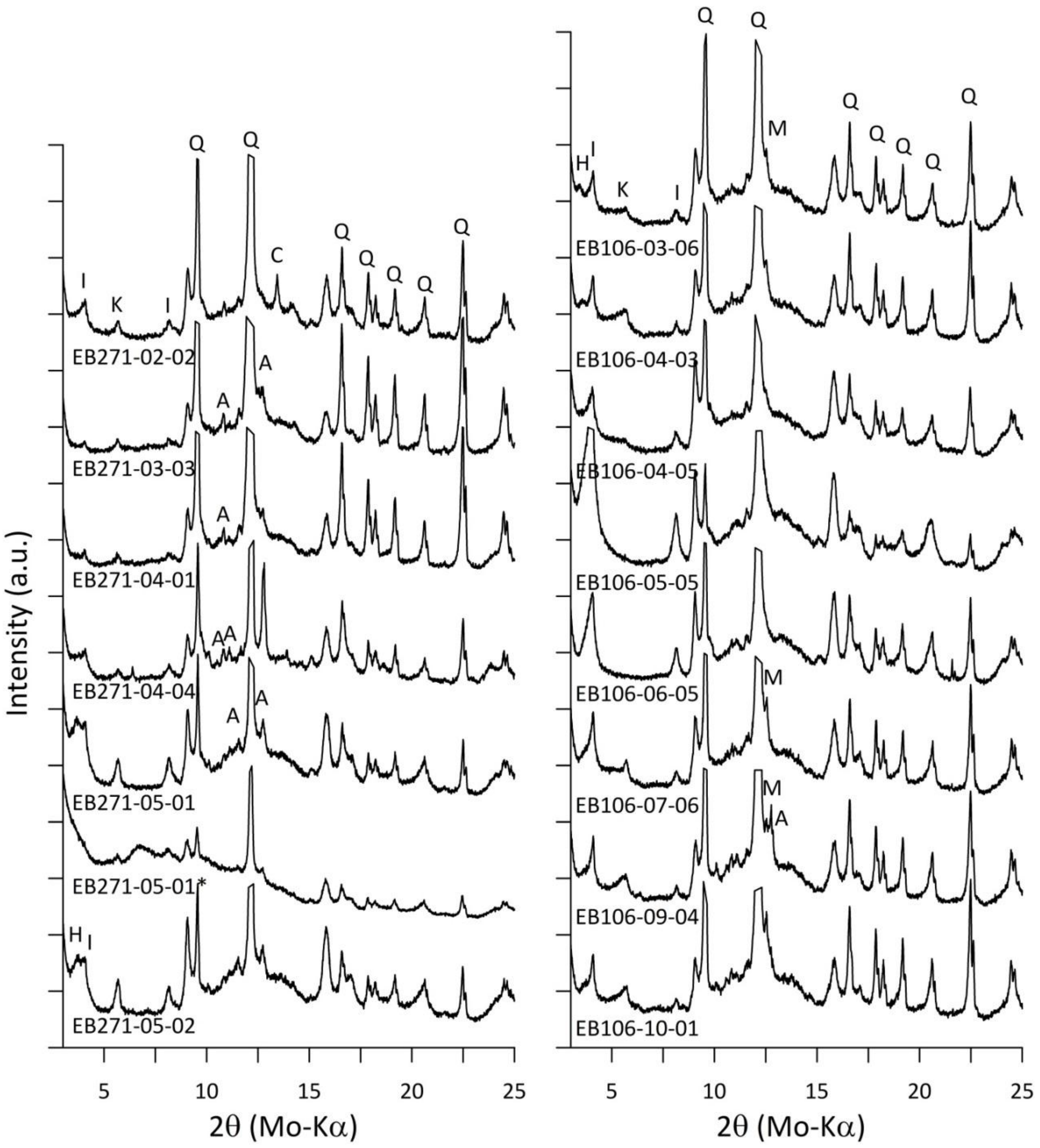


Supplemental Figure S1. The XRD patterns from selected samples from EB271 and EB106 core sediments.

Set (a)

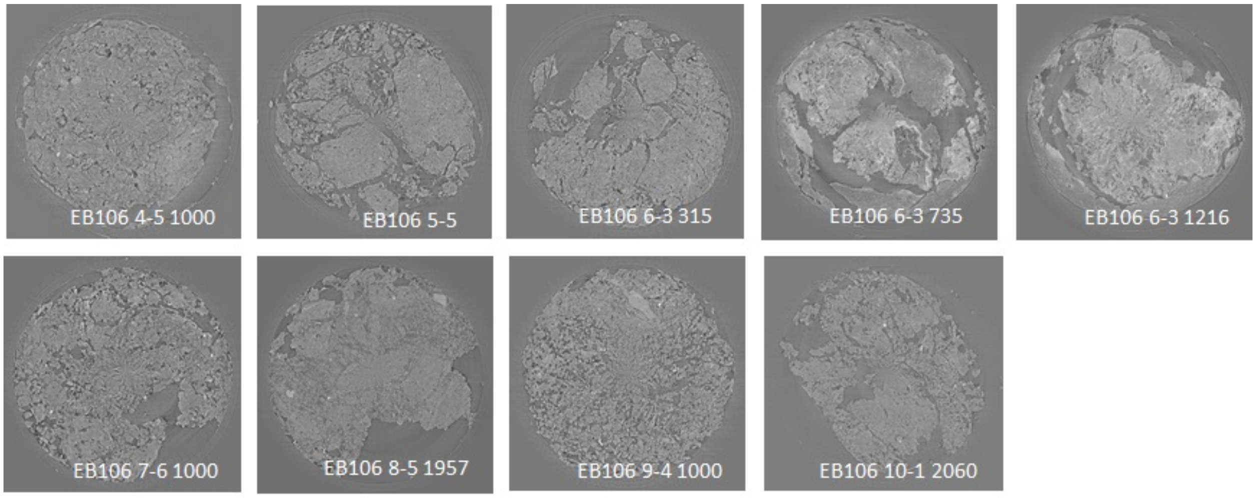


Set (b)

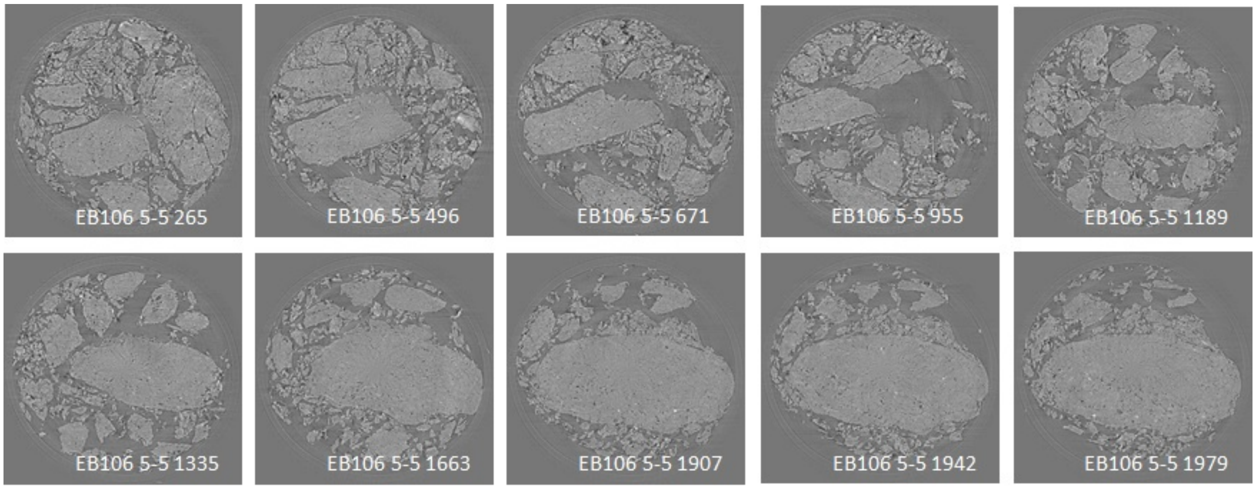


Sec (c)

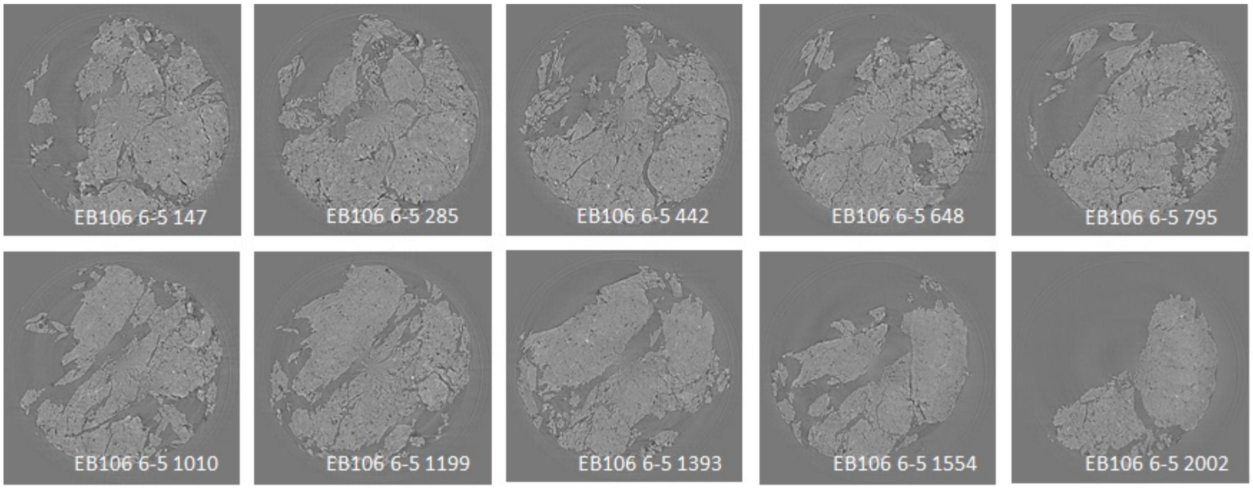

Supplemental Figure S2. The microCT images from (a) EB 106 core sediment, (b) EB106, section 5-5 (426–441 cm bgs) and (c) EB106 6-5 (527–549 cm bgs).

Supplemental Table 1. Groundwater wells GW217 and FW106, Well Construction in ft/in.

| | | |
|---|---|---|
| LOC_ID | GW271 | FW106 |
| Longitude | -84.27058705 | -84.2734838 |
| Latitude | 35.97989558 | 35.97729757 |
| LOC_ORIG_DATE | 6/5/86 | 6/9/03 |
| TOT_DRILLED_DEPTH | 56.3 | 47.08 |
| NORTHING_VAL | 30434.56 | 30228.38 |
| EASTING_VAL | 53234.33 | 51955.82 |
| TOC_ELEV | 1009.01 | 1008.53 |
| GS_ELEV | 1006.49 | 1008.12 |
| RISER_CASING_ID | 4 | 3.975 |
| RISER_CASING_BOT_DEPTH | 45.9 | 45 |
| SCREEN_TYPE | SLS/SW | SLS/SW |
| SCREEN_OPENING_SIZE | 0.01 | 0.01 |
| SCREEN_TOP_DEPTH | 45.9 | 35.25 |
| SCREEN_BOT_DEPTH | 56.3 | 45 |
| COND_CASING_TYPE | PVC40 | PVC40 |

Supplemental Table 2. GPS of boreholes EB271 and EB106.

| | | |
|---|---|---|
| LOC_ID | EB271 | EB106 |
| Longitude | -84.27059 | -84.2735 |
| Latitude | 35.97986 | 35.8772 |

**Methods**

**Inference of sediment segment depth below ground surface (bgs)**

Sediment cores were collected of approximately 91.44 cm and cut into 22.86 segments. To infer the bgs location of these segments (Supplemental Tables 3 and 4), we accounted for the deformation (expansion and compaction) of sediment cores during the coring process. We used a linear core adjustment to approximate the deformation (Morton *et al*. 2009)

$$D_{adj} = D\left(\frac{P}{L}\right) \quad (1)$$

$$D_{inf} = D_{pre} + D_{adj} \quad (2)$$

where $D_{adj}$ is the adjusted core length, $D$ is the recovered depth, $P$ is the core barrel penetration length, and $L$ is the length of the sediment core (Equation 1). $P/L$ is the recovery factor (Supplemental Tables 3 and 4). For the top and bottom of each core, the depth was inferred from the penetration of the GeoProbe. For all other depths (bottom of the top segment, top of the last segment, and top and bottom for all other segments), the inferred depth $D_{inf}$ is the sum of the previous calculated depth $D_{pre}$ and adjusted core length $D_{adj}$ (Equation 2).

**References**

Morton RA, Bernier JC, and Buster NA. (2009) Simple Methods for Evaluating Accommodation Space Formation in Coastal Wetlands. *Wetlands*. 29(3):997-1003.

Supplemental Table 3. EB271 Depth Adjustments

| | | | | | | Inferred in bgs | | Inferred ft bgs | | Inferred cm bgs | | | |
|---|---|---|---|---|---|---|---|---|---|---|---|---|---|
| Core | Official Naming | Reovery Factor (P/L) (1) | Recovered Core Length (L) (in.) | Adjusted Core Length ($D_{adj}$) (in.) | Segment Length (in.) | Top (in bgs) | Bottom (in bgs) | Top (ft bgs) | Bottom (ft bgs) | Top (cm bgs)^ | Bottom (cm bgs)^ | Middle (cm bgs) | Deformation |
| 01-01 | EB271-00 | 1.33 | 27.00 | 36.00 | 27.00 | 0.00 | 36.00 | 0.00 | 3.00 | 0.00 | 91.44 | 46 | Compaction |
| 02-01 | EB271-01 | 1.64 | | 14.73 | 9.00 | 36.00 | 50.73 | 3.00 | 4.23 | 91.44 | 128.85 | 110 | Compaction |
| 02-02 | EB271-02 | 1.64 | 22.00 | 14.73 | 9.00 | 50.73 | 65.45 | 4.23 | 5.45 | 128.85 | 166.25 | 148 | Compaction |
| 02-03 | EB271-03 | 1.64 | | 6.55 | 4.00 | 65.45 | 72.00 | 5.45 | 6.00 | 166.25 | 182.88 | 175 | Compaction |
| 03-01 | EB271-04 | 1.24 | | 11.17 | 9.00 | 72.00 | 83.17 | 6.00 | 6.93 | 182.88 | 211.26 | 197 | Compaction |
| 03-02 | EB271-05 | 1.24 | 29.00 | 11.17 | 9.00 | 83.17 | 94.34 | 6.93 | 7.86 | 211.26 | 239.64 | 225 | Compaction |
| 03-03 | EB271-06 | 1.24 | | 13.66 | 11.00 | 94.34 | 108.00 | 7.86 | 9.00 | 239.64 | 274.32 | 257 | Compaction |
| 04-01 | EB271-07 | 1.03 | | 9.26 | 9.00 | 108.00 | 117.26 | 9.00 | 9.77 | 274.32 | 297.83 | 286 | Compaction |
| 04-02 | EB271-08 | 1.03 | 35.00 | 9.26 | 9.00 | 117.26 | 126.51 | 9.77 | 10.54 | 297.83 | 321.35 | 310 | Compaction |
| 04-03 | EB271-09 | 1.03 | | 9.26 | 9.00 | 126.51 | 135.77 | 10.54 | 11.31 | 321.35 | 344.86 | 333 | Compaction |
| 04-04 | EB271-10 | 1.03 | | 8.23 | 8.00 | 135.77 | 144.00 | 11.31 | 12.00 | 344.86 | 365.76 | 355 | Compaction |
| 05-01 | EB271-11 | 0.77 | | 6.89 | 9.00 | 144.00 | 150.89 | 12.00 | 12.57 | 365.76 | 383.27 | 375 | Expansion |
| 05-02 | EB271-12 | 0.77 | | 6.89 | 9.00 | 150.89 | 157.79 | 12.57 | 13.15 | 383.27 | 400.78 | 392 | Expansion |
| 05-03 | EB271-13 | 0.77 | 47.00 | 6.89 | 9.00 | 157.79 | 164.68 | 13.15 | 13.72 | 400.78 | 418.29 | 410 | Expansion |
| 05-04 | EB271-14 | 0.77 | | 8.04 | 10.50 | 164.68 | 172.72 | 13.72 | 14.39 | 418.29 | 438.72 | 429 | Expansion |
| 05-05* | EB271-15 | 0.77 | | 7.28 | 9.50 | 172.72 | 180.00 | 14.39 | 15.00 | 438.72 | 457.20 | 448 | Expansion |
| 06-01 | EB271-16 | 0.35 | 10.00 | 3.50 | 10.00 | 180.00 | 183.50 | 15.00 | 15.29 | 457.20 | 466.09 | 462 | Expansion |

Notes

(1) Recovery Factor is the length of core advanced divided by the length of core recovered, e.g., 36-inch advance of the core from 3 to 6 feet below ground surface (ft bgs) and 24-inch recovery of the core would result in a Recovery Factor of 1.5. Recovery Factors greater than 1 indicate under-recovery. Recovery Factors less than 1 indicate over-recovery.

(*) Not processed. Stuck in bottom of barrel in field, but included for depth measurements.

### Supplemental Table 4. EB106 Depth Adjustments.

| | | | | | | Inferred in bgs | | Inferred ft bgs | | Inferred cm bgs | | | |
|---|---|---|---|---|---|---|---|---|---|---|---|---|---|
| Core-Segment | Official Naming | Reovery Factor (P/L) (1) | Recovered Core Length (L) (in.) | Adjusted Core Length ($D_{adj}$) (in.) | Segment Length (in) | Top (in bgs) | Bottom (in bgs) | Top (ft bgs) | Bottom (ft bgs) | Top (cm bgs) | Bottom (cm bgs) | Middle (cm bgs) | Deformation |
| 01-01 | EB106-00 | 1.80 | 20.00 | | 20.00 | 0.00 | 36.00 | 0.00 | 3.00 | 0.00 | 91.44 | 45.72 | Compaction |
| 02-01 | EB106-01 | 1.57 | | 14.09 | 9.00 | 36.00 | 50.09 | 3.00 | 4.17 | 91.44 | 127.22 | 109.33 | Compaction |
| 02-02 | EB106-02 | 1.57 | 23.00 | 14.09 | 9.00 | 50.09 | 64.17 | 4.17 | 5.35 | 127.22 | 163.00 | 145.11 | Compaction |
| 02-03 | EB106-03 | 1.57 | | 7.83 | 5.00 | 64.17 | 72.00 | 5.35 | 6.00 | 163.00 | 182.88 | 172.94 | Compaction |
| 03-01 | EB106-04 | 0.64 | | 5.79 | 9.00 | 72.00 | 77.79 | 6.00 | 6.48 | 182.88 | 197.58 | 190.23 | Expansion |
| 03-02 | EB106-05 | 0.64 | | 5.79 | 9.00 | 77.79 | 83.57 | 6.48 | 6.96 | 197.58 | 212.27 | 204.92 | Expansion |
| 03-03 | EB106-06 | 0.64 | 56.00 | 5.79 | 9.00 | 83.57 | 89.36 | 6.96 | 7.45 | 212.27 | 226.97 | 219.62 | Expansion |
| 03-04 | EB106-07 | 0.64 | | 5.79 | 9.00 | 89.36 | 95.14 | 7.45 | 7.93 | 226.97 | 241.66 | 234.32 | Expansion |
| 03-05 | EB106-08 | 0.64 | | 5.79 | 9.00 | 95.14 | 100.93 | 7.93 | 8.41 | 241.66 | 256.36 | 249.01 | Expansion |
| 03-06 | EB106-09 | 0.64 | | 7.07 | 11.00 | 100.93 | 108.00 | 8.41 | 9.00 | 256.36 | 274.32 | 265.34 | Expansion |
| 04-01 | EB106-10 | 0.75 | | 6.75 | 9.00 | 108.00 | 114.75 | 9.00 | 9.56 | 274.32 | 291.47 | 282.89 | Expansion |
| 04-02 | EB106-11 | 0.75 | | 6.75 | 9.00 | 114.75 | 121.50 | 9.56 | 10.13 | 291.47 | 308.61 | 300.04 | Expansion |
| 04-03 | EB106-12 | 0.75 | 48.00 | 6.75 | 9.00 | 121.50 | 128.25 | 10.13 | 10.69 | 308.61 | 325.76 | 317.18 | Expansion |
| 04-04 | EB106-13 | 0.75 | | 6.75 | 9.00 | 128.25 | 135.00 | 10.69 | 11.25 | 325.76 | 342.90 | 334.33 | Expansion |
| 04-05 | EB106-14 | 0.75 | | 9.00 | 12.00 | 135.00 | 144.00 | 11.25 | 12.00 | 342.90 | 365.76 | 354.33 | Expansion |
| 05-01 | EB106-15 | 0.65 | | 5.89 | 9.00 | 144.00 | 149.89 | 12.00 | 12.49 | 365.76 | 380.72 | 373.24 | Expansion |
| 05-02 | EB106-16 | 0.65 | | 5.89 | 9.00 | 149.89 | 155.78 | 12.49 | 12.98 | 380.72 | 395.69 | 388.20 | Expansion |
| 05-03 | EB106-17 | 0.65 | 55.00 | 5.89 | 9.00 | 155.78 | 161.67 | 12.98 | 13.47 | 395.69 | 410.65 | 403.17 | Expansion |
| 05-04 | EB106-18 | 0.65 | | 5.89 | 9.00 | 161.67 | 167.56 | 13.47 | 13.96 | 410.65 | 425.61 | 418.13 | Expansion |
| 05-05 | EB106-19 | 0.65 | | 5.89 | 9.00 | 167.56 | 173.45 | 13.96 | 14.45 | 425.61 | 440.57 | 433.09 | Expansion |
| 05-06 | EB106-20 | 0.65 | | 6.55 | 10.00 | 173.45 | 180.00 | 14.45 | 15.00 | 440.57 | 457.20 | 448.89 | Expansion |
| 06-01 | EB106-21 | 0.77 | | 6.89 | 9.00 | 180.00 | 186.89 | 15.00 | 15.57 | 457.20 | 474.71 | 465.95 | Expansion |
| 06-02 | EB106-22 | 0.77 | | 6.89 | 9.00 | 186.89 | 193.79 | 15.57 | 16.15 | 474.71 | 492.22 | 483.46 | Expansion |
| 06-03 | EB106-23 | 0.77 | 47.00 | 6.89 | 9.00 | 193.79 | 200.68 | 16.15 | 16.72 | 492.22 | 509.73 | 500.97 | Expansion |
| 06-04 | EB106-24 | 0.77 | | 6.89 | 9.00 | 200.68 | 207.57 | 16.72 | 17.30 | 509.73 | 527.24 | 518.48 | Expansion |
| 06-05 | EB106-25 | 0.77 | | 8.43 | 11.00 | 207.57 | 216.00 | 17.30 | 18.00 | 527.24 | 548.64 | 537.94 | Expansion |
| 07-01 | EB106-26 | 0.60 | | 5.40 | 9.00 | 216.00 | 221.40 | 18.00 | 18.45 | 548.64 | 562.36 | 555.50 | Expansion |
| 07-02 | EB106-27 | 0.60 | | 5.40 | 9.00 | 221.40 | 226.80 | 18.45 | 18.90 | 562.36 | 576.07 | 569.21 | Expansion |
| 07-03 | EB106-28 | 0.60 | 60.00 | 5.40 | 9.00 | 226.80 | 232.20 | 18.90 | 19.35 | 576.07 | 589.79 | 582.93 | Expansion |
| 07-04 | EB106-29 | 0.60 | | 5.40 | 9.00 | 232.20 | 237.60 | 19.35 | 19.80 | 589.79 | 603.50 | 596.65 | Expansion |
| 07-05 | EB106-30 | 0.60 | | 5.40 | 9.00 | 237.60 | 243.00 | 19.80 | 20.25 | 603.50 | 617.22 | 610.36 | Expansion |
| 07-06 | EB106-31 | 0.60 | | 9.00 | 15.00 | 243.00 | 252.00 | 20.25 | 21.00 | 617.22 | 640.08 | 628.65 | Expansion |
| 08-01 | EB106-32 | 0.60 | | 5.40 | 9.00 | 252.00 | 257.40 | 21.00 | 21.45 | 640.08 | 653.80 | 646.94 | Expansion |
| 08-02 | EB106-33 | 0.60 | | 5.40 | 9.00 | 257.40 | 262.80 | 21.45 | 21.90 | 653.80 | 667.51 | 660.65 | Expansion |
| 08-03 | EB106-34 | 0.60 | 60.00 | 5.40 | 9.00 | 262.80 | 268.20 | 21.90 | 22.35 | 667.51 | 681.23 | 674.37 | Expansion |
| 08-04 | EB106-35 | 0.60 | | 5.40 | 9.00 | 268.20 | 273.60 | 22.35 | 22.80 | 681.23 | 694.94 | 688.09 | Expansion |
| 08-05 | EB106-36 | 0.60 | | 5.40 | 9.00 | 273.60 | 279.00 | 22.80 | 23.25 | 694.94 | 708.66 | 701.80 | Expansion |
| 08-06 | EB106-37 | 0.60 | | 9.00 | 15.00 | 279.00 | 288.00 | 23.25 | 24.00 | 708.66 | 731.52 | 720.09 | Expansion |
| 09-01 | EB106-38 | 0.63 | | 5.68 | 9.00 | 288.00 | 293.68 | 24.00 | 24.47 | 731.52 | 745.96 | 738.74 | Expansion |
| 09-02 | EB106-39 | 0.63 | 38.00 | 5.68 | 9.00 | 293.68 | 299.37 | 24.47 | 24.95 | 745.96 | 760.40 | 753.18 | Expansion |
| 09-03 | EB106-40 | 0.63 | | 5.68 | 9.00 | 299.37 | 305.05 | 24.95 | 25.42 | 760.40 | 774.83 | 767.61 | Expansion |
| 09-04 | EB106-41 | 0.63 | | 6.95 | 11.00 | 305.05 | 312.00 | 25.42 | 26.00 | 774.83 | 792.48 | 783.66 | Expansion |
| 10-01 | EB106-42 | 0.72 | 12.50 | 9.00 | 12.50 | 312.00 | 321.00 | 26.00 | 26.75 | 792.48 | 815.34 | 803.91 | Expansion |

Notes

(1) Recovery Factor is the length of core advanced divided by the length of core recovered, e.g., 36-inch advance of the core from 3 to 6 feet below ground surface (ft bgs) and 24-inch recovery of the core would result in a Recovery Factor of 1.5. Recovery Factors greater than 1 indicate under-recovery. Recovery Factors less than 1 indicate over-recovery.

(*) Indicates the middle of the vertical interval of a fully homogenized core section, e.g., a core advanced from 12 to 15 ft bgs with no under-recovery would have a depth-integrated mid-point or middle of 13.5 ft bgs
